# A canonical excitatory–inhibitory motif embedded in the connectome underlies rhythmic locomotion in *Drosophila* larvae

**DOI:** 10.64898/2026.08.31.748252

**Authors:** Takaki Seki, Hiroshi Kohsaka, Maarten F Zwart, Akinao Nose

**Affiliations:** Department of Physics, Graduate School of Science, The University of Tokyo, Tokyo, Japan; Department of Complexity Science and Engineering, Graduate School of Frontier Sciences, The University of Tokyo, Chiba, Japan; Graduate School of Informatics and Engineering, University of Electro-Communications, Tokyo, Japan; School of Psychology and Neuroscience, Centre of Biophotonics, University of St Andrews, St Andrews, United Kingdom

## Abstract

Rhythmic locomotion is generated by central pattern generators, yet how defined neural circuits generate rhythmic motor waves remains unclear. Theoretical models, including Wilson–Cowan-type excitatory–inhibitory networks, have long proposed that reciprocal interactions between excitatory and inhibitory neuronal populations can generate oscillatory activity and travelling waves, but whether such motifs are physically embedded within biological connectomes is unknown. Here we show that *Drosophila* larval forward locomotion is generated by a segmentally repeated excitatory–inhibitory network centred on the excitatory A18f and inhibitory A02j neurons. Connectomic analysis revealed that A18f and A02j form reciprocally connected loops coupled across segments, receive input from forward command systems and project to premotor modules controlling muscle contraction and relaxation. Connectome-constrained firing-rate simulations, in which synaptic weights were determined by reconstructed synapse numbers, reproduced rhythmic forward-propagating motor waves with minimal parameter tuning. Physiological recordings revealed activity dynamics and phase relationships consistent with model predictions. Acute and chronic perturbations showed that A18f is required for forward locomotion, whereas optogenetic activation of posterior A18f neurons initiated forward motor waves. Perturbation of inhibitory input to A18f further supported a role for A02j-mediated inhibition. These findings identify a canonical excitatory–inhibitory motif embedded within the connectome as a biological implementation of a rhythm-generating network.

## Introduction

Rhythmic movements such as walking, swimming, breathing and peristalsis are fundamental motor behaviours shared across the animal kingdom. These movements are thought to be generated by central pattern generators (CPGs), neural circuits capable of producing rhythmic motor activity even in the absence of patterned sensory inputs (Grillner and Wallén, 1985; Marder and Calabrese, 1996; Marder and Bucher, 2001; Grillner, 2006; Friesen and Kristan, 2007; Katz, 2016). Although many neurons involved in locomotor coordination, such as left-right and flexor-extensor coordination, have been identified in vertebrate and invertebrate systems, the core circuit mechanisms that generate rhythmic activity remain incompletely understood, particularly in large nervous systems (Zhen and Samuel, 2015; Kiehn, 2016; Dougherty, 2023; Büschges and Ache, 2024; Gosgnach, 2025; Goulding et al., 2025; El Manira, 2026).

Theoretical studies have provided a complementary framework for understanding how rhythmic activity can arise from neural circuits. Early population-level models showed how recurrent interactions among neuron-like elements and excitatory–inhibitory populations can give rise to collective neural dynamics, including oscillatory activity (Amari, 1971, 1972; Wilson and Cowan, 1972). Spatially extended neural-field models and coupled-oscillator frameworks further showed how local rhythmic dynamics can give rise to travelling waves (Amari, 1977; Ermentrout and Kleinfeld, 2001; Harris and Ermentrout, 2018). However, despite the conceptual importance of these models, direct biological implementations of such rhythm-generating motifs within identified locomotor circuits remain poorly defined.

One major obstacle has been the difficulty of combining comprehensive circuit architecture with causal genetic access to identified neurons within larger, segmentally organized locomotor systems. Recent electron microscopy (EM)-based reconstructions of the *Drosophila* nervous system now provide an opportunity to bridge this gap. By revealing synaptic connectivity among identified neurons at cellular resolution in a genetically accessible animal, these connectomes make it possible to relate circuit architecture to neuronal function and behaviour (Winding, 2023; Scheffer et al., 2020; Takemura et al., 2024; Azevedo et al., 2024; Dorkenwald et al., 2024). However, it remains largely unknown which identified neurons and synaptic architectures within biological connectomes implement theoretically proposed rhythm-generating dynamics.

*Drosophila* larval crawling offers a particularly tractable context in which to address this question (Gowda et al., 2021; Kohsaka, 2023). During forward locomotion, rhythmic waves of neural activity propagate from posterior to anterior segments in the ventral nerve cord and drive corresponding waves of muscle contraction across the body (Fig. 1a; Heckscher et al., 2012; Pulver et al., 2015; Zarin et al., 2019). This system therefore provides an opportunity to ask how synaptic circuit architecture gives rise to rhythmic waves of neural activity and, ultimately, to behaviour. Previous modelling studies have proposed circuit mechanisms for the generation and propagation of peristaltic motor waves in crawling larvae (Gjorgjieva et al., 2013; Pehlevan et al., 2016; Loveless and Webb, 2018; Zarin et al., 2019; Francis et al., 2025). In parallel, experimental studies have identified premotor interneurons involved in sequential segmental contraction and relaxation (Zwart et al., 2016; Fushiki et al., 2016; Kohsaka et al., 2019; Hiramoto et al., 2021), left–right coordination (Heckscher et al., 2015), and speed control (Kohsaka et al., 2014; Liu et al., 2023; Zhao et al., 2024). These studies established important components of the larval locomotor circuit, particularly those that coordinate motor output across segments and body sides. Nevertheless, the identified neurons and synaptic architecture responsible for generating the rhythmic motor waves themselves have remained unclear.

**Figure 1.**
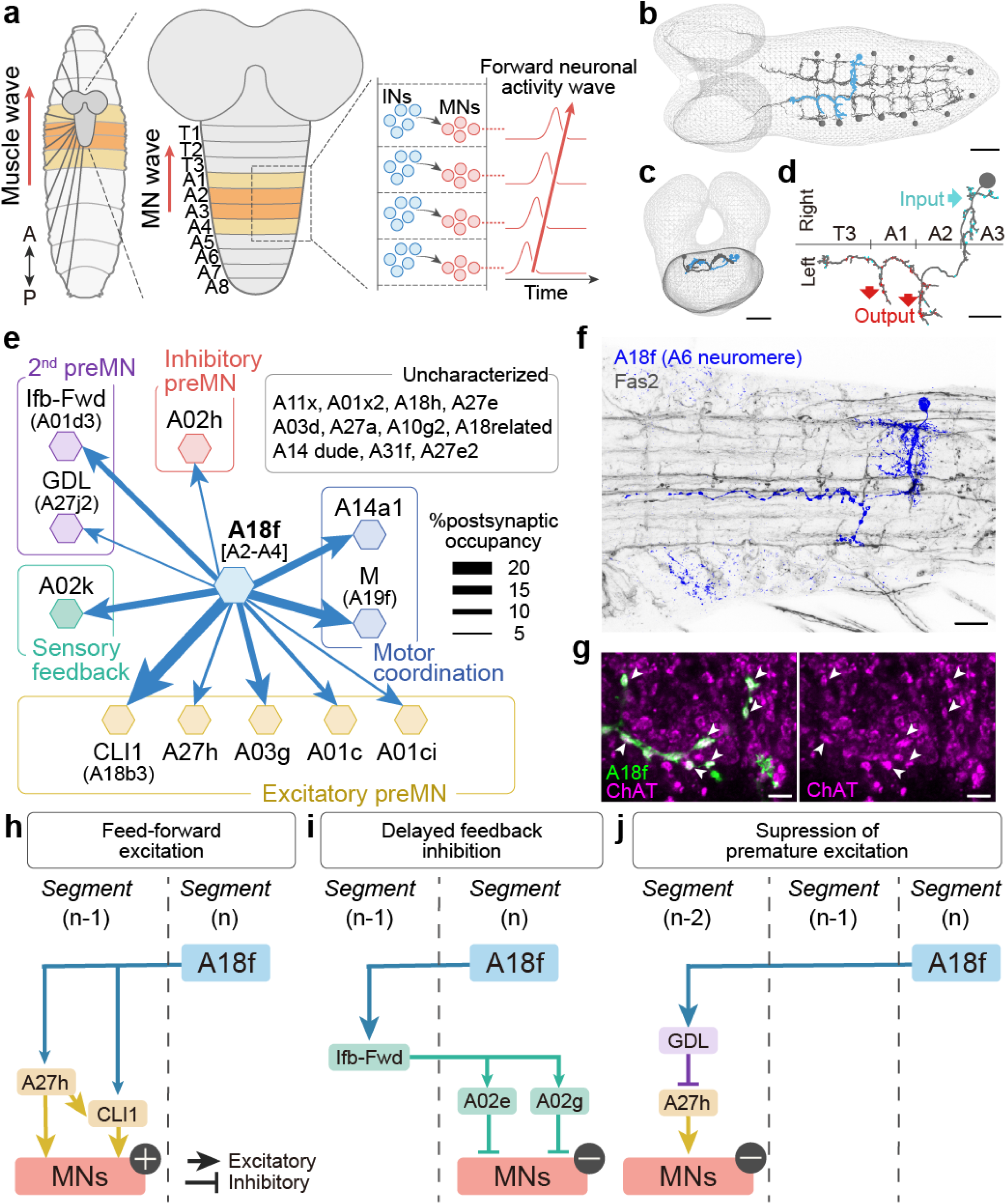
Anatomy and connectivity of A18f neurons. **a**, Schematic of larval forward locomotion and the underlying neural activity. Left, the larval body is segmented, and each body-wall segment is innervated by motor neurons located in the corresponding neuromere in the ventral nerve cord (VNC). During forward locomotion, muscle contractions propagate from posterior to anterior segments as a peristaltic wave. This motor pattern is generated by corresponding waves of neuronal activity in the VNC. Active body-wall and VNC segments are indicated in dark yellow. Right, schematic activity traces showing sequential activation of VNC segments during forward wave propagation. A, anterior; P, posterior; IN, interneuron; MN, motor neuron. **b–d**, EM reconstruction of A18f neurons. **b**, Dorsal view of A18f neurons in abdominal neuromeres A1-A8. The A18f neuron in the right A3 hemi-neuromere is highlighted in blue; grey circles indicate cell bodies. **c**, Posterior view of bilateral A18f neurons in A3. **d**, Dorsal view of the A18f neuron in the right A3 hemi-neuromere. Blue and red puncta indicate input and output sites, respectively. Scale bar, 20 µm (b, c), 10 µm (d). **e**, Downstream targets of A18f neurons identified from connectome data (A2–A4). Only neurons with postsynaptic occupancy greater than 5% are shown. Arrow thickness is proportional to postsynaptic occupancy. **f**, Confocal image of a single A18f neuron labelled by MCFO (blue). Fas2 staining was used as a neuropil landmark and shown in grey. Scale bar, 20 µm. **g**, Choline acetyltransferase (ChAT; magenta) signal is detected at A18f axon terminals labelled by *A18f-spGal4 > UAS-mCD8::GFP* (green). Arrowheads indicate ChAT-positive A18f boutons. Scale bar, 5 µm. **h–j**, Connectivity motifs between A18f and downstream premotor neurons inferred from connectome data. **h**, Feed- forward excitation. A18f innervates the excitatory premotor neuron A27h and CLI1 in the adjacent anterior neuromere, promoting motor neuron activation in that neuromere. **i**, Delayed feedback inhibition. A18f innervates the second-order premotor neuron Ifb-Fwd in the adjacent anterior segment, which recruits inhibitory premotor neurons, including A02e and A02g, one segment posteriorly to suppress motor neuron activity. **j**, Suppression of premature excitation. A18f innervates GDL two segments anteriorly, which inhibits A27h and suppress motor activation. Arrows and T-bars indicate excitatory and inhibitory connections, respectively.

Here, by combining connectomics, connectome-constrained simulations, physiological recordings, and genetic perturbations, we identify a segmentally repeated excitatory–inhibitory circuit centered on A18f and A02j interneurons that functions upstream of previously identified premotor circuits. We show that this circuit forms an intersegmentally coupled E–I network whose connectivity architecture closely resembles canonical rhythm-generating motifs proposed in theoretical models.

Simulations constrained by synapse numbers extracted from the connectome robustly reproduce propagating motor waves, and physiological recordings reveal activity dynamics consistent with these predictions. Finally, genetic perturbations demonstrate that this circuit is necessary and sufficient for rhythmic forward locomotion. Together, these findings identify a canonical excitatory–inhibitory motif embedded within the connectome that functions as a biological implementation of a rhythm-generating network.

## Results

### A18f is a common upstream neuron implicated in forward crawling

To identify neurons that may coordinate rhythmic motor wave propagation, we searched the CNS reconstruction from a first-instar larval serial-section transmission electron microscopy (ssTEM) dataset (Schneider-Mizell et al., 2016; Winding et al., 2023) for higher-order interneurons upstream of previously identified premotor neurons preferentially associated with forward locomotion: A27h, CLI1/A18b3, and Ifb-fwd/A01d3. A27h and CLI1 are excitatory premotor interneurons that innervate motor neurons within the same neuromere (Fushiki et al., 2016; Hasegawa et al., 2016). Ifb-fwd neurons are second-order premotor interneurons that inhibit motor neurons in the posterior neuromere through inhibitory premotor interneurons, including A02e and A02g (Kohsaka et al., 2019). Although these neurons have been proposed to coordinate sequential activation and inhibition of motor neurons along the body axis, they are unlikely to fully account for motor wave generation on their own. We therefore hypothesized that upstream neurons exist that orchestrate the timely propagation of activity among these premotor circuits.

This search identified A18f neurons, segmentally repeated bilateral pairs of interneurons in the ventral nerve cord, as the strongest common upstream partners of A27h, CLI1, and Ifb-fwd neurons (Fig. 1b, c; Extended Data Fig. 1a). A18f neurons also target multiple premotor interneuron classes in addition to A27h, CLI1, and Ifb-fwd, including A03g and GDL, which have previously been implicated in motor control (Fig. 1e; Supplementary Table 1; Fushiki et al., 2016; Kohsaka et al., 2019). The arrangement of A18f input and output sites was consistent with a role in feed-forward activation during forward locomotion: A18f neurons receive synaptic inputs through dendrites located near the cell body and send outputs to anterior neuromeres through axons spanning multiple neuromeres (Fig. 1d).

To determine the sign of these synaptic connections, we identified a Gal4 line that sparsely labels A18f neurons (*A18f-spGal4*; Fig. 1f; Extended Data Fig. 1b) and examined their neurotransmitter phenotype. A18f neurons expressed a cholinergic marker, but not GABAergic or glutamatergic markers, suggesting that they are excitatory neurons (Fig. 1g; Extended Data Fig. 1c, d).

Together, the EM-derived connectivity and excitatory neurotransmitter phenotype of A18f neurons suggest that A18f neurons are positioned to coordinate the timely activation and inactivation of motor neurons during forward locomotion as follows (Fig. 1h–j). First, by activating A27h and other excitatory premotor interneurons such as CLI1 in the next anterior neuromere, A18f neurons could mediate feed-forward motor activation (Fig. 1h). Second, by activating Ifb-fwd neurons in the next anterior neuromere, A18f neurons may contribute to delayed suppression of motor activity in the same neuromere through downstream inhibitory premotor interneurons (Fig. 1i). Third, by activating GDL neurons in the second anterior neuromere, A18f neurons may suppress premature motor activation in more anterior segments (Fig. 1j). Thus, A18f neurons occupy a circuit position from which they could coordinate forward motor wave propagation through multiple premotor pathways.

### A18f and A02j neurons form an intersegmental excitatory-inhibitory network downstream of forward command systems

Connectomic analysis of neurons upstream of A18f in the A1 neuromere identified A02j as one of its strongest synaptic partners (Extended Data Fig. 2a; Supplementary Table 2). A02j neurons are paired inhibitory interneurons present in each abdominal neuromere (Kohsaka et al., 2019; Zhao et al., 2024; Extended data Fig. 2c, d). Notably, this connectivity is reciprocal: A02j neurons are major upstream partners of A18f neurons, and A18f neurons are major upstream partners of A02j neurons (Extended Data Fig. 2b; Supplementary Table 3). Thus, A18f and A02j form reciprocally connected excitatory–inhibitory (E–I) loops within each neuromere (Fig. 2a).

**Figure 2.**
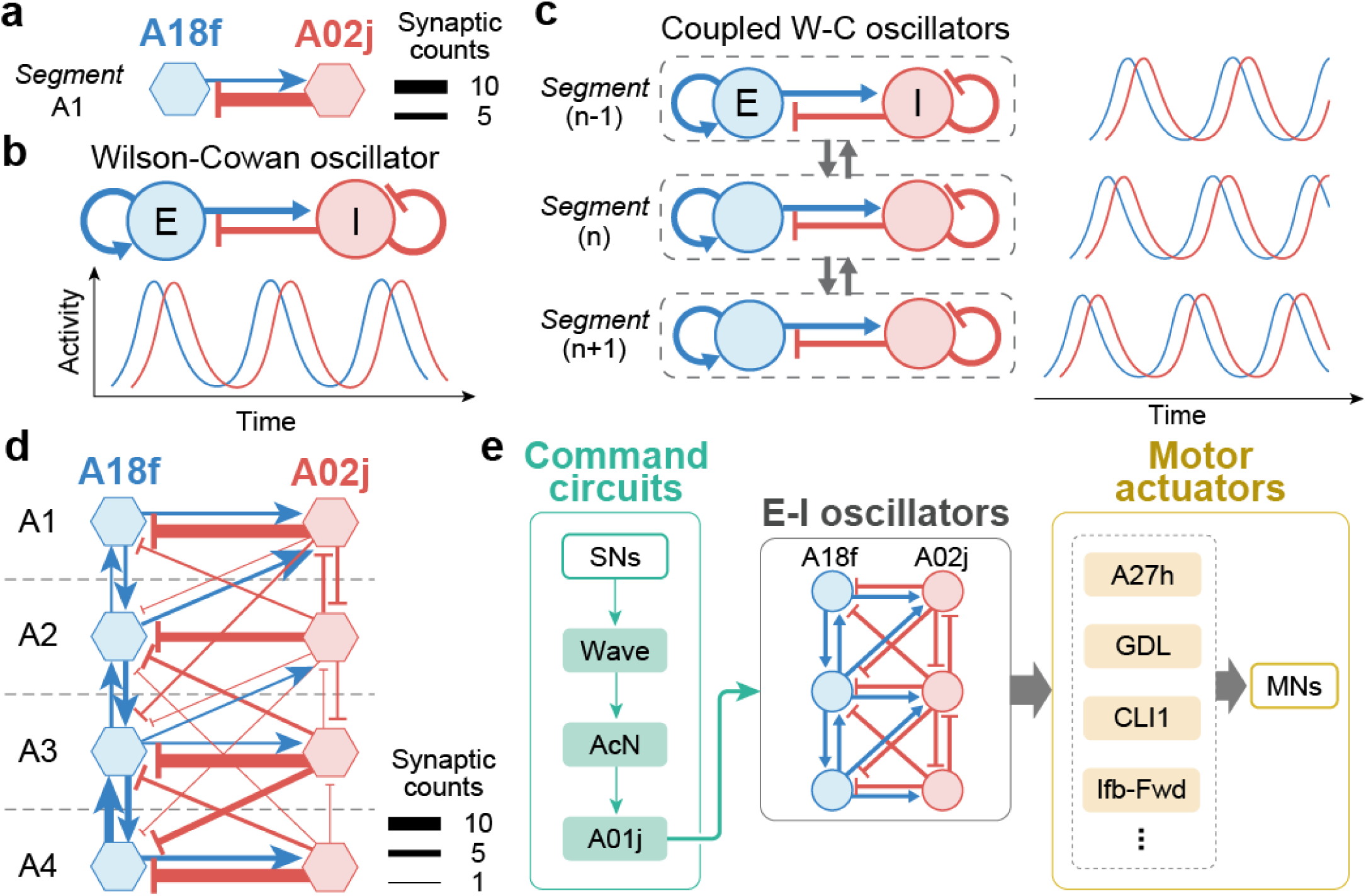
A18f and A02j neurons form a Wilson-Cowan-like oscillator network. **a**, A18f-A02j connectivity diagram in A1 based on connectome data. Left and right homologous neurons are pooled into a single node. Arrow thickness indicates synaptic count. **b**, Schematic of a Wilson-Cowan-type excitatory-inhibitory oscillator. Reciprocal interactions between an excitatory node (blue) and an inhibitory node (red) can generate oscillatory activity under appropriate conditions. **c**, Schematic of coupled excitatory-inhibitory oscillators. Coupling local oscillators across segments can generate propagating activity waves with phase delays between neighboring segments.. **d**, Connectome-derived A18f-A02j circuit diagram in A1–A4 neuromeres, showing a Wilson–Cowan-like arrangement of reciprocal excitatory and inhibitory connectivity. A18f and A02j neurons are shown in blue and red, respectively. Left and right homologous neurons within each neuromere are pooled into a single node. **e**, Schematic of the sensory-to-motor pathway for larval forward locomotion, inferred from connectome data. Command circuits for fast forward locomotion provide inputs to the A18f-A02j Wilson-Cowan-like motif, which in turn innervates premotor actuator circuits controlling motor neurons.

Such E-I motifs have long been proposed in theoretical models, including Wilson–Cowan-type models, to generate rhythmic activity (Fig. 2b; Amari, 1971, 1972; Wilson and Cowan, 1972). In these models, excitatory neurons activate inhibitory neurons, which in turn feed back to suppress excitation. Because inhibition builds up with a delay, excitation rises first and is subsequently suppressed, leading to oscillatory activity. Importantly, when local oscillators are coupled across segments, theoretical studies predict that their activity becomes phase-shifted, producing traveling waves (Fig. 2c; Amari, 1977; Ermentrout and Kleinfeld, 2001; Harris and Ermentrout, 2018).

We therefore examined whether the A18f/A02j loops are interconnected across segments in the larval connectome. Indeed, A18f and A02j were interconnected not only within individual neuromere but also across segments, forming a chain of E–I units along the body axis (Fig. 2d). Notably, the circuit architecture identified in the connectome closely resembled that predicted by theoretical models of peristaltic locomotion based on Wilson-Cowan-type dynamics (Gjorgjieva et al., 2016; Extended Data Fig. 2e, f). Together, these observations raised the possibility that the A18f/A02j network represents a biological implementation of a canonical excitatory–inhibitory rhythm-generating motif.

To identify upstream inputs to the A18f/A02j network, we further analyzed neurons presynaptic to A18f and A02j in the posterior neuromeres A7 and A8, where forward peristaltic waves initiate. This analysis identified command circuits composed of Wave, AcN, and A01j neurons, which were previously shown to induce forward locomotion as an escape response to tail touch (Fig. 2e; Takagi et al., 2017; Zhao et al., 2024). Thus, the A18f/A02j E–I network occupies a central position within the locomotor hierarchy, receiving inputs from command circuits and projecting to motor actuator circuits.

### Connectome-constrained simulations of the A18f/A02j network reproduce rhythmic motor waves

Theoretical studies predict that coupled excitatory–inhibitory circuits can generate traveling waves, but it remains unclear whether biological connectomes contain sufficient connectivity structure to implement such dynamics. We therefore asked whether the connectivity architecture identified in the larval connectome could reproduce rhythmic wave propagation. To address this question, we performed simulations using a simple firing-rate model composed of four consecutive A18f/A02j excitatory–inhibitory loops (Fig. 3a). In this model, synaptic weights were constrained by the number of synapses identified in the EM reconstruction of A18f and A02j neurons in the A1–A4 neuromeres.

**Figure 3.**
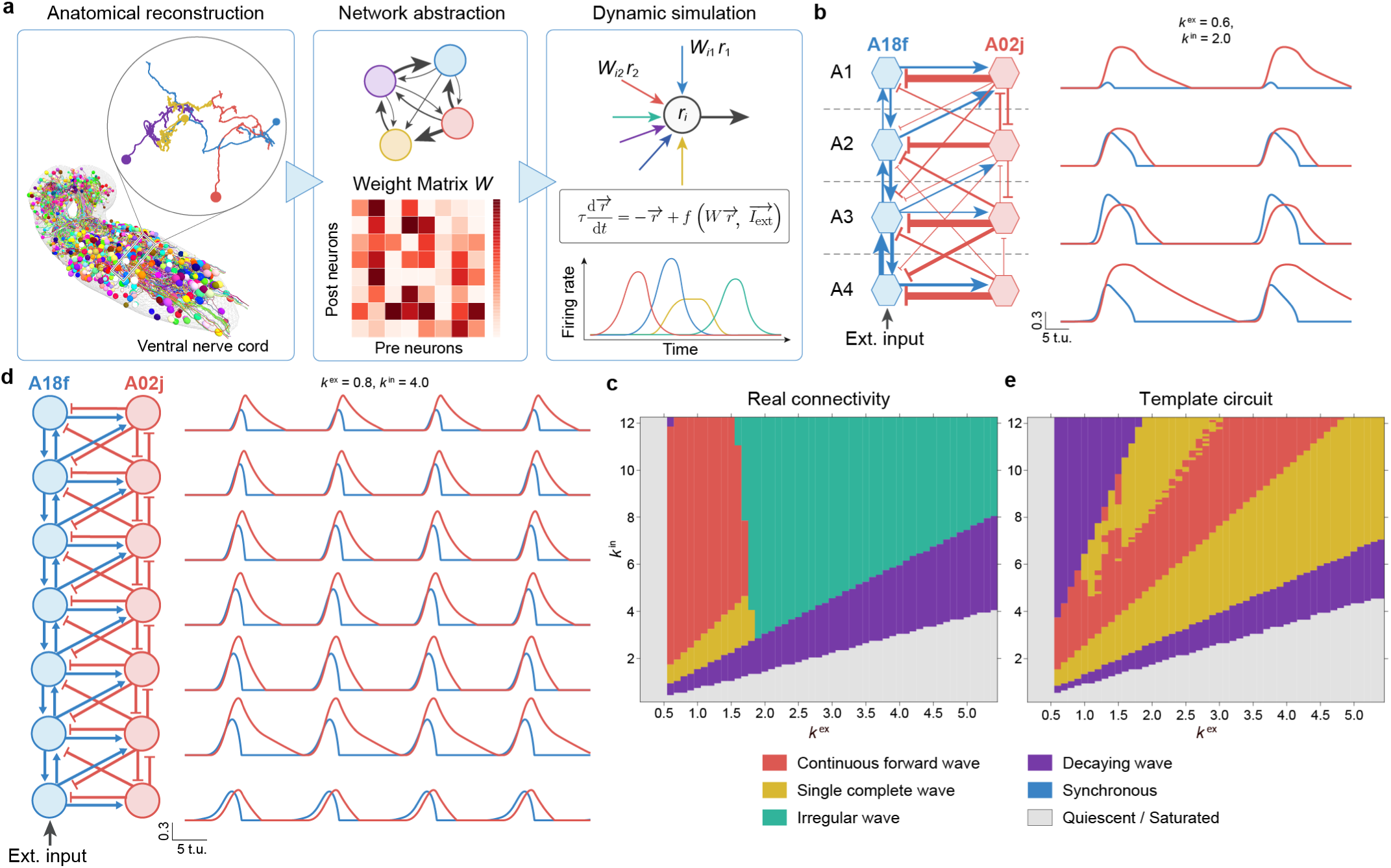
Connectome-constrained modeling and simulation of the A18f-A02j circuit. **a**, Schematic of the simulation pipeline. Neurons of interest were reconstructed from electron microscopy (EM) images to extract a connectivity graph and a synaptic weight matrix, *W*. Neuronal firing rates were simulated by incorporating this connectome-derived weight matrix into a firing rate model. **b**, Simulation of the connectome-derived A18f-A02j circuit in A1-A4 neuromeres. The circuit model corresponds to that shown in Fig. 2d and was based on EM-derived synapse counts, with left and right homologous neurons pooled into single nodes. A constant external input was applied to the A18f node in A4. Simulated activity traces show rhythmic forward-propagating waves generated by the model. *k*^ex^ = 0.6 , *k*^in^ = 2.0, external input amplitude = 2.0. **c**, Parameter space analysis of the simulation shown in **b**. Simulated waveforms obtained across different values of *k*^ex^ and *k*^in^ were classified into six categories and are shown as a phase diagram: continuous forward waves (red), single complete waves (orange), irregular waves (green), decaying waves (purple), synchronized activity (blue), and quiescent or saturated activity, in which activity either did not exceed the detection threshold or remained elevated after reaching a peak (gray). Irregular waves were defined as waveforms with insufficient intersegmental coordination, including variable activity timing across segments. Decaying waves were defined as waveforms in which activity peaks decreased substantially during anterior propagation. See Methods for detailed phase definitions. **d**, Simulation of a templated A18f/A02j circuit extended across seven neuromeres. The circuit model was generated by extending an average A18f/A02j connectivity template. Simulated activity traces show stable rhythmic forward-propagating waves. *k*^ex^ = 0.8 , *k*^in^ = 4.0, external input amplitude = 2.0. **e**, Parameter space analysis of the simulation shown in **d**, using the same classification criteria as in **c**.

When continuous external input was applied to the posterior-most A18f neurons, the model generated forward-propagating waves (Fig. 3b). Notably, although the external input to posterior A18f neurons was constant, the simulated A18f/A02j circuit autonomously generated rhythmic traveling waves: as one wave reached the anterior end, a new wave emerged from the posterior end. Wave generation was observed over broad ranges of model parameters, including the strength of tonic input to the posterior-most A18f neurons and the global excitatory and inhibitory scaling factors (*k*^ex^ and *k*^in^), provided that excitation–inhibition balance was maintained (Fig. 3c and data not shown). Thus, rhythmic propagation emerged from a connectivity architecture constrained by connectome-derived synapse numbers, without tuning individual synaptic weights or fitting dynamical parameters to locomotor kinematics.

Although the four-neuromere simulation reproduced rhythmic waves, activity in the terminal neuromeres, A1 and A4, was somewhat irregular, potentially because the modeled circuit was artificially truncated from more anterior and posterior neuromeres.

Because complete reconstruction of the A18f/A02j network across all abdominal neuromeres is currently technically challenging, we generated a template A18f/A02j circuit using the average number of synapses observed across A1–A4 and extended this template to model an A1–A7 circuit. Simulation of this extended A18f/A02j circuit generated more regular rhythmic traveling waves in a robust manner (Fig. 3d, e). These findings suggest that the A18f/A02j connectivity identified in the connectome is sufficient, in a simple firing-rate framework, to generate rhythmic traveling waves characteristic of larval forward locomotion.

We also examined the minimal structural requirements for rhythmic activity by simulating truncated forms of the template A18f/A02j circuit. In many excitatory–inhibitory oscillator models, recurrent excitation contributes to the stabilization of sustained oscillations. However, the larval connectome reveals that A18f neurons lack recurrent self-connections. Consistent with this anatomical constraint, simulation of an isolated single-segment A18f–A02j circuit failed to sustain rhythmicity, producing only decaying oscillations (Extended Data Fig. 3a, b). Strikingly, however, coupling just two or three segments was sufficient to stabilize the dynamics and autonomously generate rhythmic propagating activity (Extended Data Fig. 3c–f). Linear stability analysis provided a possible explanation for this effect, suggesting that intersegmental excitatory coupling can act as an effective recurrent-excitation-like term at the network level (Supplementary Note). Thus, the template A18f/A02j circuit cannot sustain oscillation as a single unit, but can generate stable wave-like activity when coupled across segments.

We next examined whether both excitatory A18f neurons and inhibitory A02j neurons are required for wave generation in silico. Although A18f neurons in neighboring neuromeres form an intersegmentally coupled excitatory chain, the A18f- only circuit failed to generate propagating waves (Extended Data Fig. 4a, b). Likewise, the A02j-only circuit was also insufficient to generate rhythmic propagation (Extended Data Fig. 4c, d). Thus, excitatory–inhibitory interactions between A18f and A02j neurons were required for robust wave generation in the model. We also examined whether another class of inhibitory neurons, A02k, which we found in the connectome to form reciprocal connection with A18f, could function like A02j. Although A18f and A02k also form a chain of excitatory-inhibitory loops along the neuromeres, they fail to form waves in the simulation (Extended Data Fig. 4e, f). Thus, wave generation requires not only reciprocal excitatory–inhibitory connectivity, but also the specific intersegmental connectivity pattern of the A18f/A02j network.

### The A18f/A02j network exhibits activity dynamics consistent with connectome-constrained simulations

We next asked whether the A18f/A02j circuit is activated during locomotion in a manner predicted by the connectome-constrained simulations. To monitor A18f activity, we used *A18f-spGAL4*, which primarily drives expression in A18f neurons in the A6 and A7 neuromeres, to express the calcium indicator GCaMP6f. To simultaneously monitor motor activity, we also expressed GCaMP6f in a subset of motor neurons (aCC and RP2), and performed calcium imaging in isolated central nervous system preparations. We found that A18f neurons were selectively active during fictive forward locomotion but remained largely inactive during backward waves (Fig. 4a). Cross-correlation analysis revealed that A18f activation occurred approximately synchronously with motor neuron activation in the next anterior neuromere (Fig. 4b–d), consistent with connectomic observations that A18f neurons innervate excitatory premotor interneurons, including A27h, in the next anterior neuromere. This forward-specific activity was further confirmed in intact larvae using the calcium-modulated photoactivatable integrator CaMPARI (Fig. 4e).

**Figure 4.**
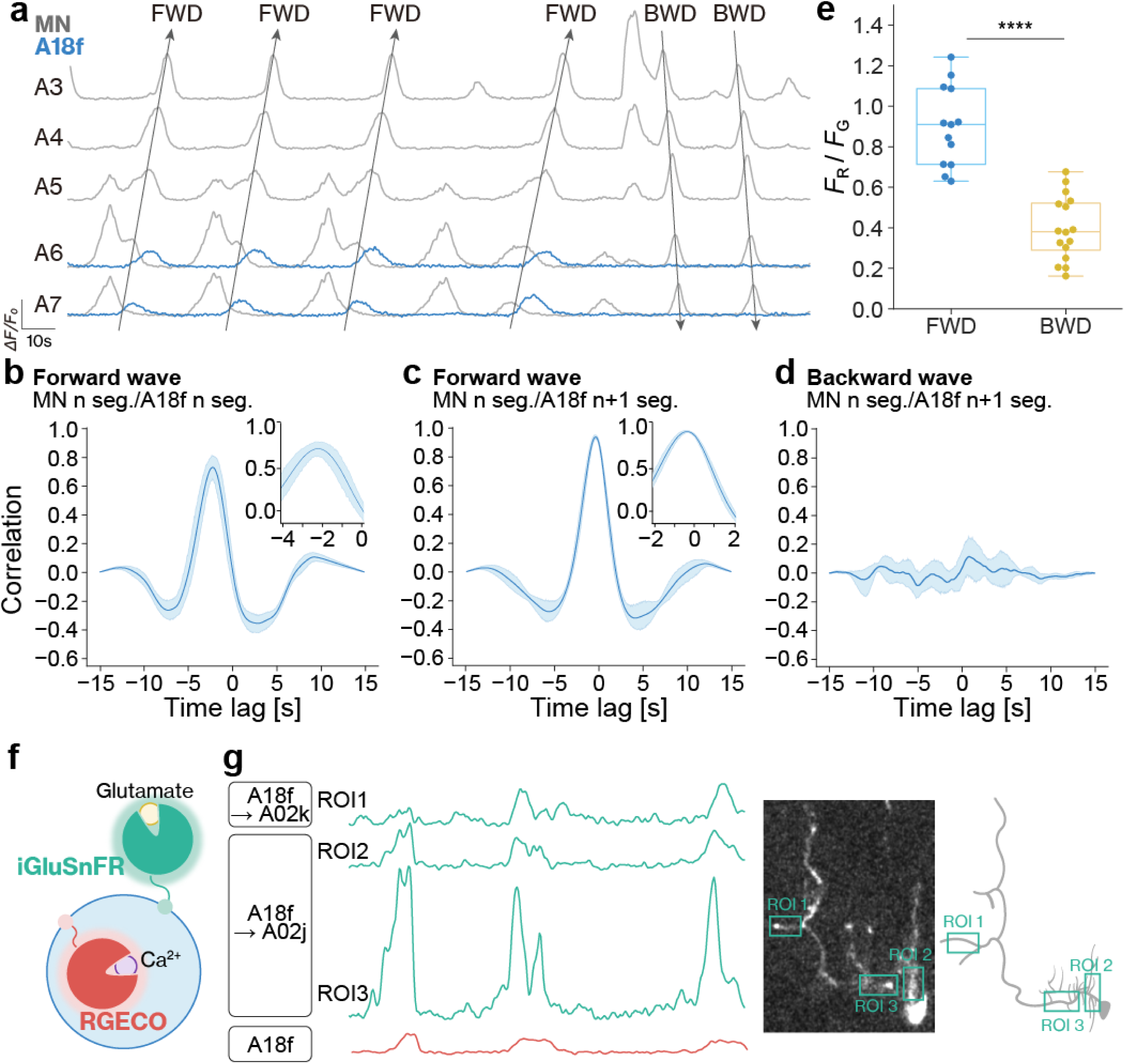
A18f neurons are preferentially active during forward locomotion and receive concurrent glutamatergic inputs. **a–d**, A18f neurons are active during forward, but not backward, fictive locomotor waves. A18f and motor neuron activity were monitored by simultaneous calcium imaging in the isolated CNS using *A18f-spGal4 > UAS- CD4::GCaMP6f* and *eve-Gal4 > UAS-CD4::GCaMP6f*, respectively. **a**, Representative activity traces of A18f neurons in A6 and A7 (blue) and motor neurons in A3–A7 (grey) during fictive locomotion. Forward (FWD) and backward (BWD) waves are indicated. **b–d**, Cross-correlation analysis of calcium transients between A18f and motor neurons during fictive locomotion. Correlations are shown between A18f and motor neurons in the same segment during forward waves (**b**), between A18f and motor neurons in the adjacent anterior neuromere during forward waves (**c**), and A18f and motor neurons in the same neuromere during backward waves (**d**). Shaded regions represent 95% confidence intervals. In all panels, *n* = 10. **e**, CaMPARI2 imaging showing preferential activation of A18f neurons during forward crawling in intact larvae. Quantification of the photoconversion of A18f neurons during forward crawling (blue, *n* = 13 from 6 animals) and backward crawling (orange, *n* = 16 from 5 animals) in *A18f-spGal4 > UAS-CaMPARI2* larvae. \*\*\*\**P* < 0.0001, Welch’s t-test. **f, g**, Simultaneous imaging of glutamatergic input to A18f neurons and A18f calcium activity. **f**, Schematic of the experimental design. iGluSnFR3 and jRGECO1b were co-expressed in A18f neurons. iGluSnFR3 reports glutamatergic input, whereas jRGECO1b reports intracellular Ca^2+^ levels in A18f neurons. **g**, Left: Representative fluorescence traces of iGluSnFR3 from three regions of interest (ROIs) corresponding to putative A02j and A02k input sites (green), together with jRGECO1b fluorescence from the A18f neuron (red). Right, ROIs overlaid on the fluorescence image and corresponding neuronal morphology schematic.

Sparse GAL4 lines selectively labeling A02j neurons are not currently available. We therefore used *A18f-spGAL4* to express the glutamate sensor iGluSnFR3 in A18f neurons and monitored glutamatergic inputs onto these neurons, part of which reflects A02j activity. One complication is that another glutamatergic interneuron, A02k, also innervates A18f neurons as mentioned above. However, the synaptic input sites from A02j and A02k were sufficiently segregated to allow independent monitoring of glutamatergic dynamics (Extended Data Fig. 5). Consistent with the simulations, simultaneous iGluSnFR3 and jRGECO1b recordings revealed glutamate release onto A18f neurons synchronized with A18f activation during the locomotor cycle (Fig. 4f, g).

Together, these observations indicate that the A18f/A02j network exhibits activity dynamics consistent with those predicted by the connectome-constrained model during forward locomotion.

### The A18f/A02j network is necessary for rhythmic forward locomotion

We performed genetic perturbation experiments to test whether the A18f/A02j network is required for forward locomotion in larvae. To acutely inhibit A18f neurons, we used *A18f-spGAL4* to express the light-gated chloride channel GtACR1 and optogenetically silenced these neurons during locomotion. Upon light exposure, larvae exhibited a robust suppression of forward crawling (Fig. 5a-d; Supplementary Video 1). As described above, *A18f-spGAL4* drives stochastic expression in subsets of A18f neurons in the A6 and A7 neuromeres. We therefore dissected and immunostained larvae following optogenetic experiments to determine the number of A18f neurons expressing GtACR1. Remarkably, forward locomotion was strongly suppressed irrespective of the number of labeled A18f neurons (Extended Data Fig. 6a). In some individuals, silencing of even a single A18f neuron was sufficient to strongly impair forward crawling, suggesting that A18f neurons play a critical role in wave generation.

**Figure 5.**
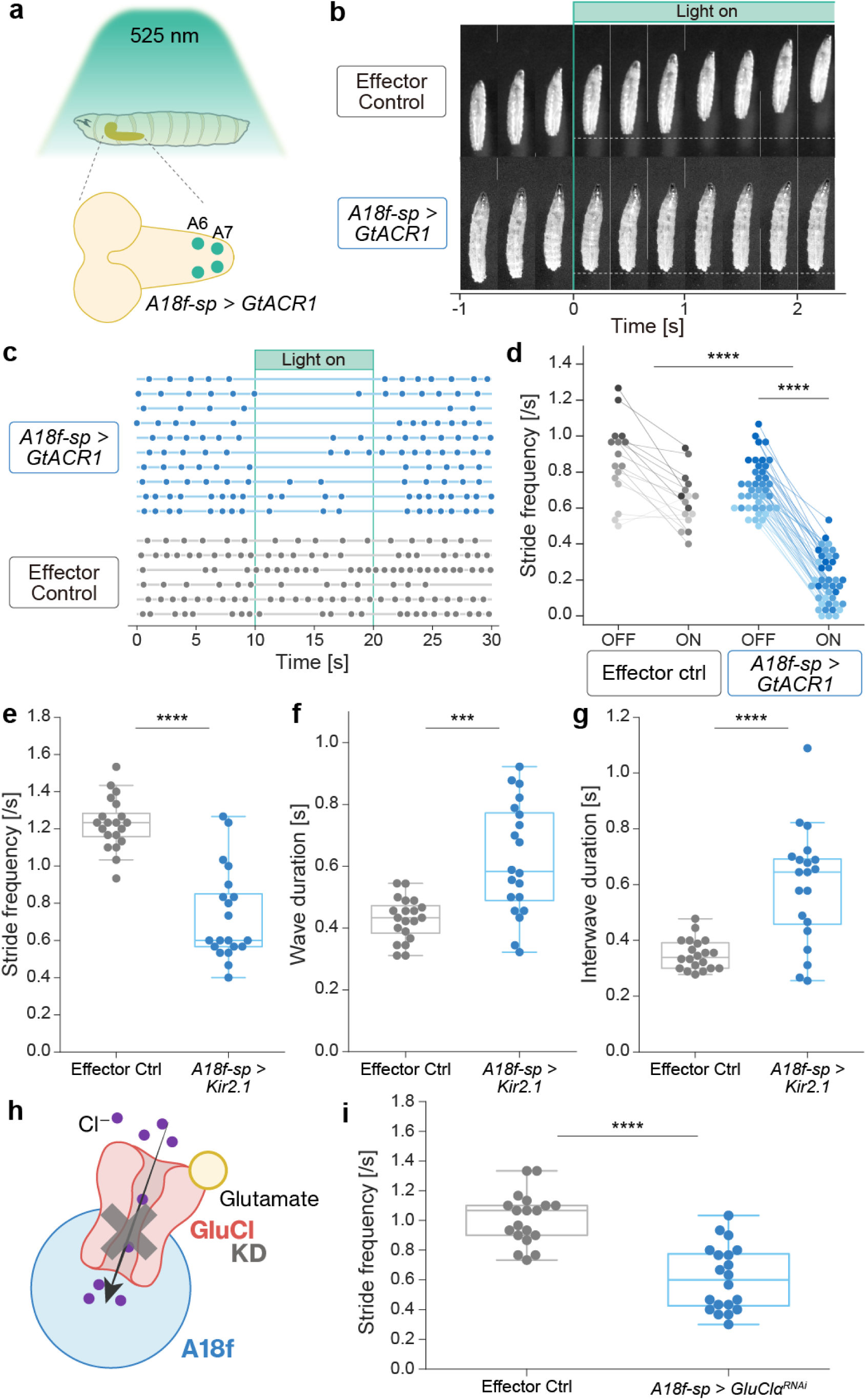
Silencing A18f neurons or reducing glutamatergic input to A18f disrupts forward locomotion in larvae. **a**, Schematic of the optogenetic silencing experiments. Larvae expressing GtACR1 in A18f neurons (*A18f-spGal4 > UAS-GtACR1*) were irradiated with green light (∼540 nm). **b–d**, Transient silencing of A18f neurons strongly impaired forward locomotion. Experimental group, *A18f-spGal4 > UAS-GtACR1*; Effector control, *UAS-GtACR1/+*. **b**, Time-lapse images of larval locomotion. Top, effector control; bottom: experimental group. Light onset is set to 0 s. **c**, Raster plot of larval forward locomotion. Each row represents a single larva, and each dot indicates a single forward crawl. **d**, Quantification of forward locomotion before and during green-light illumination. Green light induced a mild avoidance response in effector controls (grey, *n* = 16), but caused a significantly stronger reduction in forward locomotion in the experimental group (blue, *n* = 41). \*\*\*\**P* < 0.0001, paired *t*-test and Welch’s *t*-test with Holm-Bonferroni correction. **e–g**, Chronic silencing of A18f neurons by expression of the inward-rectifying potassium channel Kir2.1 impaired forward locomotion (blue, *A18f-spGal4 > UAS-Kir2.1*, *n* = 20; grey, *UAS-Kir2.1/+*, *n* = 20). Quantified parameters are forward stride frequency (**e**), wave duration defined as the time required for a muscle contraction wave to propagate from the posterior to the anterior end (**f**), and interwave duration defined as the time from an arrival of one wave at the anterior end to initiation of the next wave at the posterior end (**g**). \*\*\**P* < 0.001, \*\*\*\**P* < 0.001, Welch’s *t*-test. **h**, Schematic of the *GluClα* knockdown in A18f neurons. In *Drosophila*, the glutamate-gated chloride channel is encoded by *GluClα*. **i**, Knockdown of *GluClα* in A18f neurons reduced forward stride frequency. Experimental group, *A18f-spGal4 > UAS- GluClα^RNAi^* (blue, *n* = 20); effector control, *UAS-GluClα^RNAi^/+* (grey, *n* = 20). \*\*\*\**P* < 0.001, Welch’s t-test.

To examine the effect of chronic silencing, we next expressed the inwardly rectifying potassium channel Kir2.1 in A18f neurons. Chronic inhibition of A18f neurons severely disrupted forward locomotion (Fig. 5e; Supplementary Video 2).

Detailed analysis revealed that both wave duration and inter-wave duration were significantly prolonged (Fig. 5f, g), indicating that A18f inhibition affects not only wave propagation but also the initiation of subsequent waves. These findings suggest that A18f neurons in posterior segments play important roles in both the initiation and propagation of forward motor programs. Consistent with these behavioral phenotypes, inhibition of A18f neurons also disrupted forward wave generation during fictive locomotion in isolated central nervous system and fillet preparations (Extended Data Fig. 6b-e).

In addition to A18f neurons, A18f-spGAL4 labels a pair of off-target neurons in the subesophageal zone (SEZ). These neurons peripherally to the ring gland and/or pharyngeal muscles, and are therefore unlikely to directly regulate abdominal motor circuits. Indeed, several observations indicate that these neurons do not substantially contribute to the observed phenotypes. First, the off-target neurons were not detectably active during fictive locomotion in isolated central nervous system preparations (Extended Data Fig. 6f). Second, optogenetic inhibition continued to strongly suppress forward locomotion in fillet preparations in which the SEZ including the off-target neurons had been surgically removed (Extended Data Fig. 6d, e). Together, these observations indicate that the locomotor phenotypes produced by GtACR1-mediated inhibition primarily result from perturbation of A18f neurons.

We next asked whether A02j-mediated inhibitory feedback is also required for forward locomotion. Because GAL4 lines sparsely targeting A02j neurons are not currently available, we instead attempted to perturb glutamatergic inhibitory inputs onto A18f neurons. In *Drosophila*, inhibitory glutamatergic transmission is mediated by the glutamate-gated chloride channel GluClα (Fig. 5h). Using a *GluClα-T2A-GAL4* knock-in line, we confirmed that *GluClα* is endogenously expressed in A18f neurons (Extended Data Fig. 6g). We therefore selectively knocked down GluClα in A18f neurons using RNAi and found that this manipulation significantly reduced the frequency of forward waves (Fig. 5i). We also used *per-GAL4*, which labels a subset of glutamatergic neurons derived from the A02 lineage, including A02j but not A02k neurons, to optogenetically inhibit these neurons. Larval crawling was strongly disrupted upon photoinhibition (Extended Data Fig. 6h), consistent with a role for A02j neurons in rhythmic forward locomotion. Together, these observations suggest that both excitatory output from A18f neurons and inhibitory glutamatergic feedback onto A18f neurons are required for robust forward locomotion, with A02j neurons likely providing an important component of this feedback.

### Activation of A18f neurons induces forward locomotion

We next tested whether activation of A18f neurons is sufficient to induce forward locomotion. To this end, we used A18f-spGAL4 to express CsChrimson in A18f neurons in the posterior A6 and A7 neuromeres and optogenetically activated these neurons in larvae in a resting state. Remarkably, transient optogenetic activation of posterior A18f neurons for 1 s was sufficient to elicit forward locomotor wave(s) (Fig. 6a-c, Supplementary Video 3).

**Figure 6.**
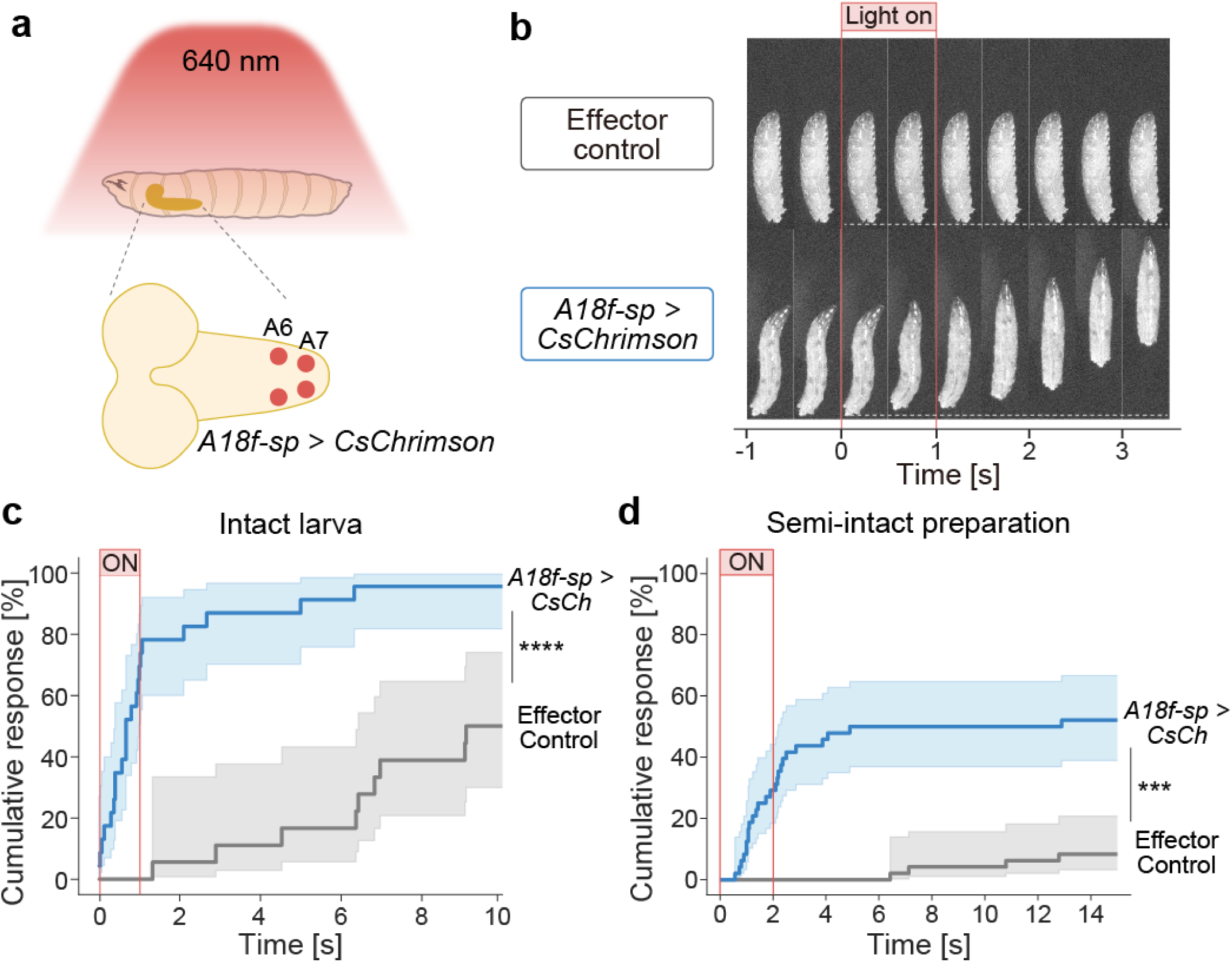
Activation of A18f neurons initiates forward motor waves. **a-c**, Brief activation of A18f neurons using the light-gated cation channel CsChrimson induced forward locomotion in resting larvae. **a**, Schematic of the optogenetic activation experiments. Larvae expressing CsChrimson in A18f neurons were stimulated with red light. **b**, Representative time-lapse images of larval locomotion. Top, effector control, *UAS-CsChrimson/+*; bottom, experimental group, *A18f-spGal4 > UAS-CsChrimson*. Light onset is set to 0 s. **c**, Cumulative incidence of forward crawling following light onset. Solid lines and shaded regions represent Kaplan-Meier estimates and 95% confidence intervals, respectively. Effector control, grey, n = 18 trials from 6 animals; experimental group, blue, *n* = 20 trials from 7 animals. \*\*\*\**P* < 0.0001, Cox proportional hazards model with cluster-robust standard errors. **d**, Cumulative incidence of forward waves following light onset in semi-intact preparations, plotted as in **c**. Effector control, grey, *n* = 48 trials from 16 animals; experimental group, blue, *n* = 48 trials from 16 animals. \*\*\**P* < 0.001, Cox proportional hazards model with cluster-robust standard errors.

We also performed optogenetic activation in fillet preparations in which dissected larvae were pinned onto a silicone substrate. Under these conditions, fictive forward locomotion gradually ceased and the preparation entered a resting state.

Transient optogenetic activation of posterior A18f neurons for 2 s was sufficient to elicit forward locomotor waves (Fig. 6d; Supplementary Video 4). Importantly, forward locomotion could still be induced after surgical removal of the SEZ, excluding contributions from off-target neurons labeled by A18f-spGAL4 (Extended Data Fig. 7).

Together, these observations indicate that activation of posterior A18f neurons is sufficient to initiate rhythmic forward locomotion.

## Discussion

In this study, we identified a segmentally repeated excitatory–inhibitory network centred on A18f and A02j interneurons that links forward command pathways to premotor circuits controlling larval crawling. Its architecture and dynamics resemble theoretical excitatory–inhibitory motifs for rhythm generation, in which delayed inhibitory feedback supports oscillation and intersegmental coupling converts local rhythms into propagating waves. Thus, the A18f/A02j network provides evidence that the larval connectome contains a biological implementation of a canonical rhythm-generating principle. This conclusion does not rest on reciprocal excitatory–inhibitory connectivity alone, as such loops may occur widely in neural circuits. Rather, the significance of the A18f/A02j network lies in the convergence of several features: its position between command pathways and premotor circuits, its segmentally repeated and intersegmentally coupled architecture, its ability to generate waves in connectome-constrained simulations, and its causal role in forward locomotion. The failure of the related A18f/A02k reciprocal loop to generate propagating waves further indicates that specific intersegmental connectivity, rather than E–I reciprocity per se, is critical.

The organization of the A18f/A02j network also provides a potential explanation for the flexibility of axial motor pattern generation. Axial motor waves have long been proposed to arise from segmentally repeated rhythm-generating units coupled along the body axis (Grillner and Wallén, 1984; Skinner and Mulloney, 1998; Friesen and Kristan, 2007). Theoretical excitatory–inhibitory models similarly show that local E–I circuits composed of neuron populations can generate rhythmic activity and, when coupled across segments, traveling waves (Amari, 1977; Harris and Ermentrout, 2018; Gjorgjieva et al., 2013). The A18f/A02j network follows this general logic, but with important distinctions: its nodes correspond to single identified neuronal classes rather than neuronal populations, and A18f neurons lack recurrent self-excitation. Accordingly, isolated single-segment A18f/A02j loops failed to sustain oscillations in the model.

Instead, rhythmic activity emerged when multiple segments were coupled, with linear stability analysis suggesting that intersegmental excitatory coupling can provide an effective recurrent-excitation-like term at the network level (Supplementary Note).

Thus, intersegmental connectivity may contribute not only to coordination between local oscillators, but also to rhythm generation itself. This modular but interdependent organization may explain how posterior tonic drive can initiate propagating waves with different numbers of coupled units, consistent with previous ablation experiments showing that fictive forward motor waves can still be generated after removal of the segments in which waves normally initiate (Pulver et al., 2015). Whether analogous coupling of local E–I modules are used more broadly in axial locomotor systems remains an important question.

We observed variability in synaptic weights among A18f/A02j loops across neuromeres and in intersegmental connections between neighboring units. This variability suggests that travelling wave generation may be supported by a range of connectivity patterns. Although the present study does not establish strict mathematical equivalence between the A18f/A02j network and classical Wilson–Cowan equations, this flexibility is consistent with Wilson–Cowan-type models of larval peristalsis, in which propagating waves arise over broad parameter regimes (Gjorgjieva et al., 2013, 2016), and with the broader principle of degeneracy in motor circuits, whereby distinct synaptic and intrinsic parameter combinations can generate similar network outputs (Prinz et al., 2004). Developmental and activity-dependent mechanisms may therefore establish A18f/A02j connectivity within a functionally permissive range rather than specify a single exact wiring pattern.

Although central circuits can generate larval locomotor waves, sensory feedback is known to shape motor output, particularly locomotor speed (Hughes et al., 2007; Song et al., 2007). A02k may provide one route by which such feedback influences the A18f-centred rhythm-generating network. Like A02j, A02k forms reciprocal connections with A18f neurons, but A02k receives stronger sensory feedback inputs (Schneider-Mizell et al., 2016). Although the A18f/A02k circuit failed to generate propagating waves in connectome-constrained simulations (Extended Data Fig. 4e, f), this does not exclude a role for A02k in wave generation within the closed sensorimotor loop, where sensory feedback could provide the intersegmental linkage that is insufficient in the central circuit alone. Thus, A02j and A02k may represent related inhibitory partners of A18f that support rhythm generation through distinct mechanisms: A02j through a central excitatory–inhibitory loop and A02k through sensorimotor feedback. Their origin as sister neurons further raises the possibility that related inhibitory interneurons have diversified to support central and feedback-dependent components of locomotor control.

The A18f/A02j network also occupies a key position within the locomotor hierarchy. Upstream, it receives input from command circuits involved in touch-evoked forward escape behaviours, including Wave, AcN, and A01j neurons. Because Wave and AcN neurons induce fast forward locomotion but are not required for normal crawling (Takagi et al., 2017; Zhao et al., 2024), additional command or modulatory pathways are likely to recruit the A18f/A02j network during spontaneous locomotion.

Downstream, A18f/A02j projects to premotor circuits implicated in motor propagation, inhibition, and intersegmental coordination. This organization suggests that the A18f/A02j network functions as a rhythm-generator through which command neurons can recruit coordinated forward locomotor programs.

Previous studies in vertebrate and invertebrate locomotor systems have identified neurons and circuit principles underlying rhythmic motor control through electrophysiology, physiological perturbation, and modelling (Robertson and Pearson, 1985; Grillner and Wallén, 1985; Friesen and Kristan, 2007; Roberts et al., 2010; Mulloney and Smarandache-Wellmann, 2012; Katz, 2016; Kiehn, 2016; Büschges and Ache, 2025). However, these systems have often relied on partial connectivity and model-based inference, whereas theoretical models have frequently been only weakly constrained by actual synaptic architecture. Our findings help bridge these levels of description by linking identified neurons, synapse-level connectivity, connectome-constrained simulations, physiological activity, and causal perturbation within a single locomotor circuit.

Recent connectome-based studies in adult flies have similarly begun to identify candidate rhythm-generating circuits through rate-coding simulations and computational pruning approaches (Pugliese et al., 2025; Sapkal et al., 2026). In the adult walking system, such analyses predicted minimal circuits capable of generating rhythmic leg motor activity in silico, although their proposed rhythm-generating roles remain to be tested experimentally. Notably, these predicted circuits also contain reciprocally connected excitatory and inhibitory neurons, resembling the A18f/A02j network identified here in larvae. Together, these observations suggest that systematic searches across connectomes may reveal recurring circuit motifs that implement shared principles of locomotor rhythm generation.

Several important questions remain. How is the A18f/A02j network assembled during development? How does it interact with sensory feedback, neuromodulatory pathways, and alternative command systems to regulate speed, behavioural state, and motor pattern selection? Are related excitatory–inhibitory motifs used to generate backward locomotion or other larval motor programs? Addressing these questions will clarify how connectome-defined circuit motifs are built, tuned, and deployed to generate flexible behaviour. Thus, connectomes may provide a route from circuit anatomy to theoretically grounded dynamical principles that generate behaviour.

## Author Contribution

T.S., M.F.Z., and A.N. conceived and designed the project and wrote the manuscript with input from the other authors. T.S. performed all experiments and analyses unless otherwise noted. H.K. initially identified A18f as an upstream neuron type of major premotor neurons. M.F.Z. performed the EM reconstruction. M.F.Z. and A.N. supervised the project.

## Acknowledgment

We are especially grateful to Dr. James Truman for generously providing the A18f split- Gal4 line. This line was indispensable for the present study, and much of this work would not have been possible without his contribution. We thank Dr. Albert Cardona for providing access to the 1st instar larval EM dataset and Dr. Kazushi Fukumasu for providing Python code for imaging analysis. We thank the Bloomington and Kyoto Stock Centers for the fly stocks. We thank Drs. Julijana Gjorgjieva, Zhefeng Gong, Qianhui Zhao, Yuri Bilk Matos, and Jimena Berni for comments on the manuscript and/or technical advice.

## Methods

### Drosophila melanogaster strains

All flies were maintained on a standard cornmeal medium at 25°C. The following strains were used in this study:

- *w^1118^* (Bloomington Drosophila Stock Center [BDSC] #3605)
- *y^1^ w^1118^* (BDSC #6598)
- *A18f-spGal4* (*VT039624-p65ADZp* in attP40 [BDSC #73089], *R75B09-ZpGDBD*
- in attP2 [BDSC # 69246])
- *R75B09-LexA* (attP40) (BDSC #61601)
- *per-Gal4* (BDSC #7127)
- *eve^RRa-F^-Gal4* (a gift from Dr. Miki Fujioka)
- *nSyb-LexA* (su(Hw)attP1) (Kyoto Stock Center #118799)
- *GluClα-T2A-Gal4* (Kyoto Stock Center #119383)
- *UAS-CD4::tdGFP* (VK33) (BDSC #35836)
- *LexAop-mCD8::GFP* (su(Hw)attP8), *UAS-mCD8::RFP* (attP18) (BDSC #32229)
- *MCFO-3* (BDSC #64087)
- *UAS-CD4::GCaMP6f* (VK5) (Kyoto Stock Center #118816)
- *LexAop-CD4::GCaMP6f* (VK5) (Kyoto Stock Center #118821)
- *UAS-CD4::jRGECO1b* (attP40) (generated in this study; see below)
- *UAS-iGluSnFR3* (VK5) (BDSC #605807)
- *UAS-CaMPARI2* (VK5) (BDSC #78316)
- *UAS-GtACR1::EYFP* (attP2) (BDSC #92983)
- *UAS-Kir2.1::EGFP* (BDSC #6596)
- *UAS-GluClα^RNAi^*(attP40) (BDSC #53356)
- *UAS-CsChrimson::mVenus* (attP40) (BDSC #55135)

### Generation of UAS-CD4::jRGECO1b (attP40) line

The UAS-CD4::jRGECO1b construct was generated as described previously (Date et al., 2026). The resulting transgene was inserted into the attP40 landing site by phiC31- mediated integration (BestGene Inc., USA) to generate UAS-CD4::jRGECO1b (attP40) transgenic fly lines.

### Immunohistochemistry

Third-instar larvae were dissected in phosphate-buffered saline (PBS) using micro- scissors. The isolated CNS preparations were mounted on adhesive glass slides (MAS-coated slide glass, S9215, Matsunami Glass, Japan). Samples were fixed in 3.7% formaldehyde for 30 min at room temperature (RT) and washed twice for 15 min each in PBS containing 0.2% Triton X-100 (PBT) at RT. After blocking in PBT containing 5% normal goat serum for 30 min at RT, the samples were incubated with primary antibodies at 4°C for overnight or longer. The samples were then washed twice for 15 min each in PBT and incubated with secondary antibodies at 4°C for overnight or longer. Following two 15-min washes in PBT, the solution was replaced with PBS.

Images were acquired using a laser scanning confocal microscope (FV3000, Evident, Japan) equipped with a 20× water-immersion objective (XLUMPLFLN 20XW, NA = 1.0, Evident, Japan), a 60× water-immersion objective (LUMPLFLN 60XW, NA = 1.0, Evident, Japan), a 100× water-immersion objective (LUMPLFL 100XW, NA = 1.0, Evident, Japan), or a 60× oil-immersion objective (UPLXAPO 60XO, NA = 1.42, Evident, Japan). For imaging with the oil-immersion objective, the sample solution was replaced with a clearing reagent (RapiClear 1.47, SunJin Lab, Taiwan) prior to observation.

The primary and secondary antibodies used, along with their dilutions, were as follows:

- Mouse anti-GFP (A-11120, Thermo Fisher Scientific), 1:1000
- Rabbit anti-GFP (GFP-Rb-Af2020, Frontier Institute), 1:1000
- Rabbit anti-DsRed (632496, Takara Bio), 1:1000
- Rabbit anti-HA (3724, Cell Signaling Technology), 1:1000
- Rat anti-FLAG (NBP1-06712, Novus Biologicals), 1:500
- Mouse anti-Fas2 (1D4, Developmental Studies Hybridoma Bank [DSHB]), 1:200
- Mouse anti-Brp (nc82, DSHB), 1:100
- Mouse anti-ChAT (4B1, DSHB), 1:50
- Rabbit anti-vGAT (Fei et al., 2010) (gift from Dr. David Krantz), 1:1000
- Rabbit anti-vGluT (Mahr & Aberle, 2006) (gift from Dr. Hermann Aberle), 1:1000
- Goat anti-mouse IgG, Alexa Fluor 488 conjugated (A-11029, Thermo Fisher Scientific), 1:300
- Goat anti-rabbit IgG, Alexa Fluor 488 conjugated (A-11034, Thermo Fisher Scientific), 1:300
- Goat anti-rat IgG, Alexa Fluor 488 conjugated (A-11006, Thermo Fisher Scientific), 1:300
- Goat anti-mouse IgG, Alexa Fluor 555 conjugated (A-21424, Thermo Fisher Scientific), 1:300
- Goat anti-mouse IgG, Alexa Fluor 647 conjugated (115-605-003, Jackson ImmunoResearch), 1:300
- Goat anti-rabbit IgG, Cy5 conjugated (A-10523, Thermo Fisher Scientific), 1:300
- Goat anti-rabbit IgG, ATTO 647N conjugated (611-156-122S, Rockland), 1:300

### Functional imaging in isolated CNS preparations

Wandering third-instar larvae were dissected in insect saline (TES buffer: 5 mM TES, 135 mM NaCl, 5 mM KCl, 4 mM MgCl_2_, 2 mM CaCl_2_, 36 mM sucrose, pH 7.15). The isolated CNS preparations were mounted on an adhesive glass slide (MAS-coated slide glass, S9215, Matsunami Glass, Japan) and bathed in the TES buffer.

For ablation experiments, after mounting on the slide, the CNS was transected around the T1-T2 segments using a single-edge razor blade so that only the VNC remained. Subsequently, the solution was replaced with TES buffer containing 1 µM pilocarpine (P6503, Merck, Germany), a muscarinic acetylcholine receptor agonist (Johnston & Levine, 1996; Ryckebusch & Laurent, 1993; Xie & Gross, 2022).

Fluorescence signals were acquired using an EMCCD camera (iXon+ DU-897E, Andor Technology, Germany) and a spinning-disk confocal unit (CSU-W1, Yokogawa, Japan) mounted on an upright microscope (BX51WI, Evident, Japan). Typically, imaging commenced immediately after mounting. In ablation experiments, however, imaging was initiated at least 8 min after ablation to reduce potential artifacts caused by injury-induced firing (Jonaitis et al., 2022).

Images were captured using either a 20× water-immersion objective (UMPLFLN 20XW, NA = 0.5, Evident, Japan) or a 40× water-immersion objective (Achroplan 40X, NA = 0.8, Zeiss, Germany). A 488 nm laser (OBIS, Coherent, USA) was used for single-color imaging, and an additional 561 nm laser (Vortran, USA) was used for dual-color imaging. Green and red fluorescence emission signals were separated by using a dual-view system (DV2, Photometrics, USA) attached to the CSU- W1 unit. Time-lapse images were acquired at a single focal plane. The acquisition rates were approximately 28 frames per second (fps) for Fig. 4a–d, 5 fps for Fig. 4g, and 8 fps for Extended Data Fig. 6b.

### Imaging data analysis

Data analysis was performed using ImageJ and custom Python 3 scripts. Minor misalignments during time-lapse imaging were corrected using the Image Stabilizer plugin in ImageJ (Li, 2008). Unless otherwise specified, regions of interest (ROIs) were manually placed over arbitrary areas of somata or neurites. Baseline detection and smoothing of fluorescence intensity traces were performed using custom Python 3 scripts based on previous studies (Eilers, 2003; Zhang et al., 2020). The detected baseline fluorescence intensity and the raw fluorescence intensity were defined as *F*_0_ and *F*, respectively. The normalized fluorescence change was calculated as Δ*F*/*F*_0_ = (*F* − *F*_0_)/*F*_0_.

For the analysis shown in Fig. 4b–d, the cross-correlation coefficient was defined as follows:

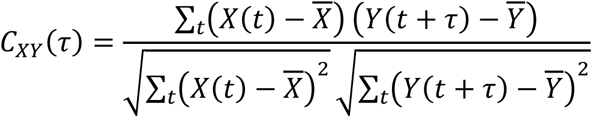

where *X*(*t*) and *Y*(*t*) are the fluorescence intensities, *X* and *Y* are their respective mean values, and *τ* is the time lag of *Y*(*t*) relative to *X*(*t*). To separate the forward- wave-related activities from posterior bursts, the fluorescence intensity of the motor neurons was fitted with a sum of two Gaussian functions using the curve_fit function in SciPy. The delayed component was considered to represent the motor neuron activity during the forward wave and was used for calculating the cross-correlation coefficient. The 95% confidence intervals for the cross-correlation coefficients were calculated using a bootstrap method with 1,000 iterations, as implemented in the lineplot function in Seaborn. All mathematical computations and visualizations were executed using Python 3 with the NumPy, SciPy, and Seaborn libraries.

### Larval locomotion assay

Wandering third-instar larvae were gently washed with deionized water, placed on a 3% agar plate containing apple juice (∼9 cm in diameter), and allowed to acclimate for at least 60 s. Larvae were then imaged for at least 60 s at 30 fps using a CCD camera (XCD-V60, Sony, Japan) mounted on a stereomicroscope (SZX16, Evident, Japan) under infrared illumination (850 nm; ∼40 µW/mm²; LDQ-150IR2-850, CCS, Japan).

Image analysis was performed using ImageJ. To quantify stride frequency, three arbitrary 10-s intervals were selected from the recording of each larva. The number of complete forward crawling strides within each interval was manually counted, and the average across the three intervals was used as the representative value for each larva. Wave duration was defined as the time from the initiation of a muscle contraction wave at the posterior end to its arrival at the anterior end. Interwave duration was defined as the time from mouth-hook unhooking to initiation of the next posterior muscle contraction wave (Liu et al., 2023). For both parameters, three forward waves were selected per larva, generally from bouts of active forward crawling, and their average was used as the representative value.

### Optogenetic assay

Second- or early third-instar larvae were transferred to agar plates spread with yeast paste containing 10 mM all-*trans*-retinal (ATR) and reared in the dark for 24–48 h.

For experiments using intact larvae, the subsequent procedures were identical to those described for the larval locomotion assay. In the GtACR1 experiments, freely crawling larvae were exposed to three 10-s pulses of green light (250 µW/mm²) separated by 15-s inter-stimulus intervals. Green light was generated using a mercury lamp (U-HGLGPS, Evident, Japan) equipped with a bandpass filter (530-550 nm; RFP1 filter, SZX2-FRFP1, Evident, Japan). In CsChrimson experiments, stationary larvae were exposed to a single 1-s pulse of red light (100 µW/mm²; 660 nm LED, M660L3, Thorlabs, USA). Imaging was continued until the larva initiated consecutive crawling movements, and three independent trials were conducted for each larva.

For experiments using semi-intact fillet preparations, larvae were washed with deionized water, transferred to a silicone-lined dish, and bathed in TES buffer. The anterior and posterior ends of the larvae were secured with insect pins. The body wall was then cut along the dorsal midline using micro-scissors and pinned flat. For experiments involving the ablation of the brain and SEZ, the CNS was subsequently transected around the T1–T2 neuromeres. For the GtACR1 experiments, only preparations that exhibited frequent forward waves post-ablation were selected.

Conversely, for the CsChrimson experiments, only preparations that remained stationary post-ablation were used. The light sources and intensities were identical to those used for intact larvae. In semi-intact GtACR1 experiments, preparations were exposed to three 10-s pulses of green light separated by 15-s intervals. In semi-intact CsChrimson experiments, preparations were exposed to three 2-s pulses of red light separated by 15- s intervals.

In the GtACR1 experiments, the number of complete forward crawling strides or waves was manually counted during the three light-on periods and during three randomly selected 10-s light-off periods. The average of these three trials were used as representative values for the light-on and light-off periods for each sample. ImageJ was used for these analyses. In the CsChrimson experiments, we quantified the latency from light onset to the first complete forward stride or wave. Survival analysis was performed, and cumulative incidence was estimated and plotted using the Kaplan-Meier method. In semi-intact preparations, trials in which no forward wave occurred within 15 s were treated as right-censored data. The 95% confidence intervals were calculated using Greenwood’s exponential formula. Analyses were performed using ImageJ and custom Python 3 scripts with the lifelines package.

### CaMPARI assay

Wandering third-instar larvae were gently washed with deionized water and placed on a 3% agar plate containing apple juice. The plate was equipped with a straight linear channel made of silicone elastomer, with a width comparable to that of a larva. Forward or backward locomotion was induced by mechanically stimulating the posterior or anterior end of a larva, respectively, using a Von Frey filament (Touch Test Sensory Evaluator [6 g], NC12775-12, North Coast Medical, USA). During induced locomotion, the larvae were illuminated with violet LED light (125 µW/mm²; 405 nm LED, M405L4, Thorlabs, USA) for 45 s. Immediately after illumination, larvae were dissected in PBS and mounted on adhesive glass slides (MAS-coated slide glass, S9215, Matsunami Glass, Japan). Samples were imaged using a confocal microscope (FV3000, Evident, Japan) equipped with a 20× water-immersion objective (XLUMPLFLN 20XW, NA = 1.0, Evident, Japan). Green and red CaMPARI fluorescence intensities were measured using 488 nm and 561 nm lasers (OBIS, Coherent, USA), respectively. The laser power was kept constant across all samples.

Image analysis was performed using ImageJ. Regions of interest (ROIs) were manually delineated over the neurites of A18f neurons to obtain mean green and red fluorescence intensities. The photoconversion level was evaluated as the red-to-green fluorescence ratio (*F_R_*/*F_G_*), calculated by dividing the mean red fluorescence intensity by the mean green fluorescence intensity.

### Statistical analysis

With the exception of survival analysis, all statistical analyses were performed using R. Statistical tests were selected based on the sample size and data distribution, from the following tests: Welch’s *t*-test, the paired *t*-test, the exact unpaired permutation test, the exact paired permutation test, and the permutation paired Brunner-Munzel test. The specific tests used for each experiment are indicated in the figure legends. The TOSTER package was used for the Brunner-Munzel test. For GtACR1 experiments, the within- subject difference between the light-on and light-off periods was calculated as the light- off value minus the light-on value to evaluate the effect of optogenetic stimulation.

For the statistical testing of survival analysis, custom Python3 scripts with the lifelines package were used. To account for within-subject correlation and avoid pseudoreplication arising from repeated trials from the same larva, a Cox proportional hazards model with cluster-robust standard errors was employed.

Statistical significance was set at an alpha level of 0.05. For multiple comparisons, *P*-values were adjusted using the Holm-Bonferroni method.

### EM connectomics analysis

EM connectomics analysis was performed using a previously published EM datasete of the CNS of a first-instar larva (Ohyama et al., 2015; Schneider-Mizell et al., 2016). For all analyses, synaptic counts from bilateral pairs of neurons were summed. To analyze the downstream targets of A18f neurons, we examined synaptic connections made by A18f neurons located in the A2–A4 neuromeres onto neurons in the A1 neuromere, where premotor neurons have been most extensively reconstructed. Downstream neurons with a postsynaptic occupancy of 5% or greater were extracted. To analyze neurons upstream of A18f and A02j, we examined neurons providing inputs to the A18f or A02j neurons in the A1 neuromere, and extracted upstream neurons with postsynaptic occupancy of 2% or greater. For the analysis of neurons upstream of A27h, Ifb-Fwd, and CLI1, we examined neurons providing input to these target neurons in the A1 neuromere. Upstream neurons were extracted if they formed synapses with at least two of the three target cell types and provided 10 or more synapses in total.

### Firing-rate simulations

To investigate the circuit dynamics using EM connectomics data, we constructed a firing-rate model based on a previous study (Jovanic et al., 2016). In this model, the

activity state of each neuron *i*, represented by its firing rate *r_i_* ≥ 0, was used as the state variable. The dynamics were defined by the following differential equation:

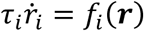

where

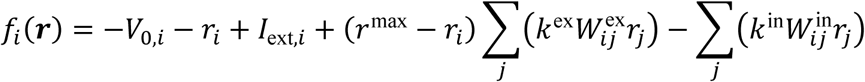

Here, *τ_i_* is the time constant, *V*_0,*i*_ is the activity threshold, *I*_ext,*i*_ is the external input, and *r*^max^ is the maximum firing rate. The element *W_ij_* of the synaptic connectivity matrix *W* denotes the number of synapses from presynaptic neuron *j* to postsynaptic neuron *i*. The excitatory connectivity matrix *W*^ex^ contains synapses from A18f neurons, while the inhibitory connectivity matrix *W*^in^ contains synapses from A02j or A02k. The parameters *k*^ex^ and *k*^in^ represent global scaling factors for excitatory and inhibitory synaptic efficacy, respectively. Connectivity matrices were generated from EM connectome by pooling left and right homologous neurons of the same cell type within each neuromere into a single node and summing their synapse numbers. The synaptic connectivity matrices used for simulations of the A18f/A02j and A18f/A02k circuits in A1-A7 neuromeres are shown in Supplementary Tables 4 and 5, respectively.

The template matrix used in the simulations was generated by averaging, across A1-A4, the synaptic inputs from each presynaptic neuron in the EM connectome, and extrapolating these averaged values across the A1-A7 neuromeres. Elements from this template matrix were utilized for simulations of truncated A18f/A02j circuits shown in Extended Data Fig. 3.

To represent external input, a single virtual neuron was added to the network. This virtual neuron had a single synaptic connection exclusively to the most posterior A18f neuron. Constant neuronal parameters, excluding those of the virtual neuron, were set as follows: *τ_i_* = 35, *V*_0,*i*_ = 20, *r*^max^ = 20, and *I*_ext,*i*_ = 0. To generate immediate and sustained activity in the virtual neuron, its parameters were set to *τ_i_* = 1, *V*_0,*i*_ = 0, *r*^max^ = 20, and *I*_ext,*i*_ = 2, with the constant external input *I*_ext,*i*_ applied continuously throughout the simulation. Numerical integration of the model dynamics was performed using the ode45 solver in MATLAB R2026a (MathWorks, Inc.), with the ’nonnegative’ option specified to ensure firing rates remained non-negative.

Based on the simulated time-series data for each neuron, we quantified wave-propagating characteristics and classified the resulting circuit dynamics into distinct phases. Wave propagation was evaluated using A18f activity in the core segments of the network, except for Extended Data Fig. 4c, for which A02j activity was analyzed. To minimize boundary effects, core segments were generally defined by excluding the anterior-most and posterior-most segments. For four-segment circuits, only the anterior- most segment was excluded, leaving three core segments. For circuits with three or fewer segments, all segments were treated as core segments.

Meaningful activity was distinguished from noise, using an activity threshold *θ* set at 1% of the maximum firing rate (*θ* = 0.01*r*^max^). Let *N_c_* denote the number of core segments. Segment indices were assigned such that the most anterior segment is *k* = 1 and the most posterior segment is *k* = *N_c_*. Using the activity of the most posterior segment (*k* = *N_c_*) as the reference, we identified firing peaks that exceeded the threshold, their peak amplitudes *A_k_*, and their onset times *t_k_* in each segment. Based on *A_k_* and *t_k_*, the following metrics were calculated to evaluate the spatial and temporal quality of wave propagation.

Amplitude Maintenance, *M*_space_, was used to evaluate whether activity amplitude was maintained during anterior propagation. It was defined as the ratio of the peak amplitude in the anterior-most core segment to that of the posterior-most core segment, capped at 1.0 :

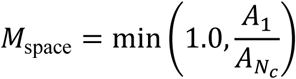

Sequence Consistency, *S*_order_, was used to evaluate whether activity propagated in the correct posterior to anterior order. In a forward wave, posterior segments fire earlier, such that *t_Nc_* < *t_Nc_*_−1_ < ⋯ < *t*_1_. A temporal penalty for retrograde propagation was defined as:

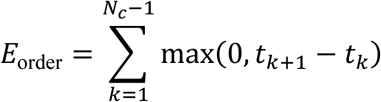

Using the total time spread, Δ*T*_total_ = max(*t_k_*) − min(*t_k_*), sequence consistency was defined as:

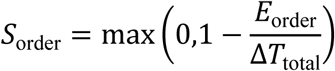

Spatial uniformity, *U*_space_, was used to evaluate whether propagation speed was uniform across segments. Inter-segment delay was defined as *d_k_* = |*t_k_*_+1_ − *t_k_*|. Using the mean *μ_d_* and standard deviation *σ_d_* of the intersegmental delays, spatial uniformity was defined as:

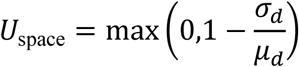

The product of these three metrics was defined as the spatial score,

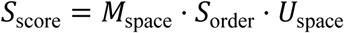

Which represents the spatial integrity of wave propagation.

The temporal score, *T*_score_, was used to evaluate whether rhythmic activity was maintained over time. A representative core segment near the centre of the simulated circuit was selected, and a list of *m* activity peak amplitudes, (*P*_1_, *P*_2_, … , *P_m_*) was detected in that segment. The temporal amplitude-maintenance score, *M*_time_, was defined as the ratio of the mean amplitude of the second and subsequent peaks to that of the first peak:

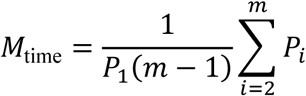

The temporal score was then defined as

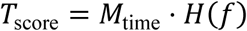

where *f* is the firing frequency estimated from the autocorrelation function and *H*(*f*) is the Heaviside step function, which returns 1 when *f* > 0 and 0 otherwise.

Based on these metrics, circuit dynamics were classified into six phases. Quiescent or saturated activity was defined as a state where no activity peaks were detected (*m* = 0), or the maximum amplitude remained below the threshold (max*_k_*(*A_k_*) < *θ*). This includes both complete silence and sustained hyperexcitable activity that rapidly saturated at a high firing rate. Synchronous activity was defined as a state in which all segments fired nearly simultaneously, with Δ*T*_total_ < 0.05. Decaying waves were defined as waves that were generated but strongly attenuated during propagation, with *M*_space_ < 0.30. Irregular waves were defined as waves in which propagation order or speed uniformity was disrupted, with *S*_order_ < 0.75 or *U*_space_ < 0.50. Single complete waves were defined as normally propagating single forward waves in which repetitive activity was not maintained, with *M*_time_ < 0.70 or *f* = 0. Continuous forward waves were defined as sustained, regular forward waves that did not meet any of the above exclusion criteria.

For single-segment A18f/A02j closed-loop simulations, spatial propagation metrics were not applied. Instead, temporal dynamics were evaluated from the A18f firing-rate, *r*(*t*), over the entire simulation period. For this analysis, the activity threshold *θ* was set to 0.001*r*^max^, and the minimum peak prominence required for oscillation was set to *θ*_osc_ = 0.005*r*^max^. Local maxima with prominence greater than or equal to *θ*_osc_ were detected as peaks with amplitudes *P*_1_, *P*_2_, … , *P_m_*. Local minima between successive peaks were denoted as troughs, *V*_1_, *V*_2_, … , *V_m_*_−1_. Oscillation persistence was quantified by the amplitude maintenance ratio, and by the final oscillation amplitude,

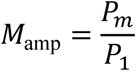

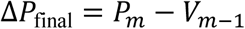

Single-segment dynamics were classified into four phases. Quiescent activity was defined as a state in which the maximum firing rate remained below the activity threshold, max(*r*(*t*)) < *θ*). Saturated activity was defined as a non-oscillatory active state in which activity exceeded the threshold but produced one or fewer peaks (*m* < 2). Damped oscillations were defined as initially oscillatory dynamics with *m* ≥ 2 that decayed over time, satisfying *M*_amp_ < 0.90 or Δ*P*_final_ < *θ*_osc_. Sustained oscillations were defined as repetitive oscillations with *m* ≥ 2 that were maintained without substantial decay, satisfying *M*_amp_ ≥ 0.90 and Δ*P*_final_ ≥ *θ*_osc_.

### Data availability

The datasets and custom code generated and analysed during the current study, including source data for all figures, are available in the figshare repository, https://doi.org/10.6084/m9.figshare.33389446.

---

**Extended data Figure 1.**
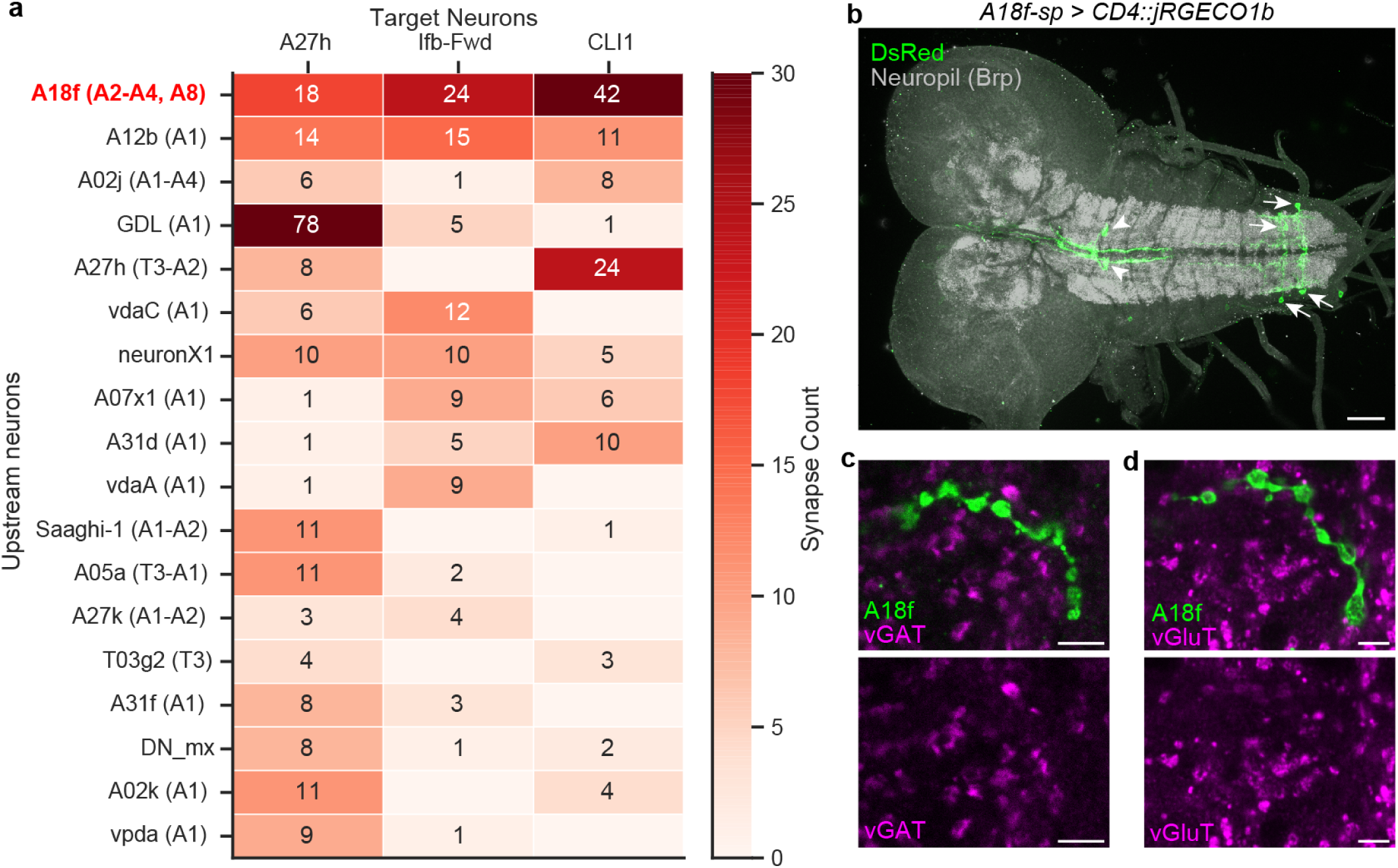
Identification and characterization of the A18f neuron. **a**, Search for common upstream partners of A27h, CLI1, and Ifb-fwd. Upstream neurons of A27h, Ifb-Fwd, and CLI1 in A1, identified from connectome data, are shown. Neurons on the y-axis are ordered first by the number of downstream target cell types they innervate and then by total synapse count. A18f was identified as the strongest common upstream partner across these forward-locomotion premotor neurons. **b**, Confocal image showing the expression pattern of *A18f-spGal4*, visualized with *A18f-spGal4 > UAS- CD4::jRGECO1b*. *A18f-spGal4* stochastically labels A18f neurons, predominantly in A6 and A7 neuromeres (arrows). It also labels a pair of unidentified cells in the subesophageal zone (SEZ) (arrowhead). Neuropil was visualized by Brp staining in grey. Scale bar, 50 µm. **c, d**, Neurotransmitter marker analysis of A18f neurons. Neither vGAT (**c**) nor vGluT (**d**) signal was detected at the axon terminals of A18f neurons labeled by *A18f-spGal4 > UAS-mCD8::GFP*. Scale bar, 5 µm.

**Extended data Figure 2.**
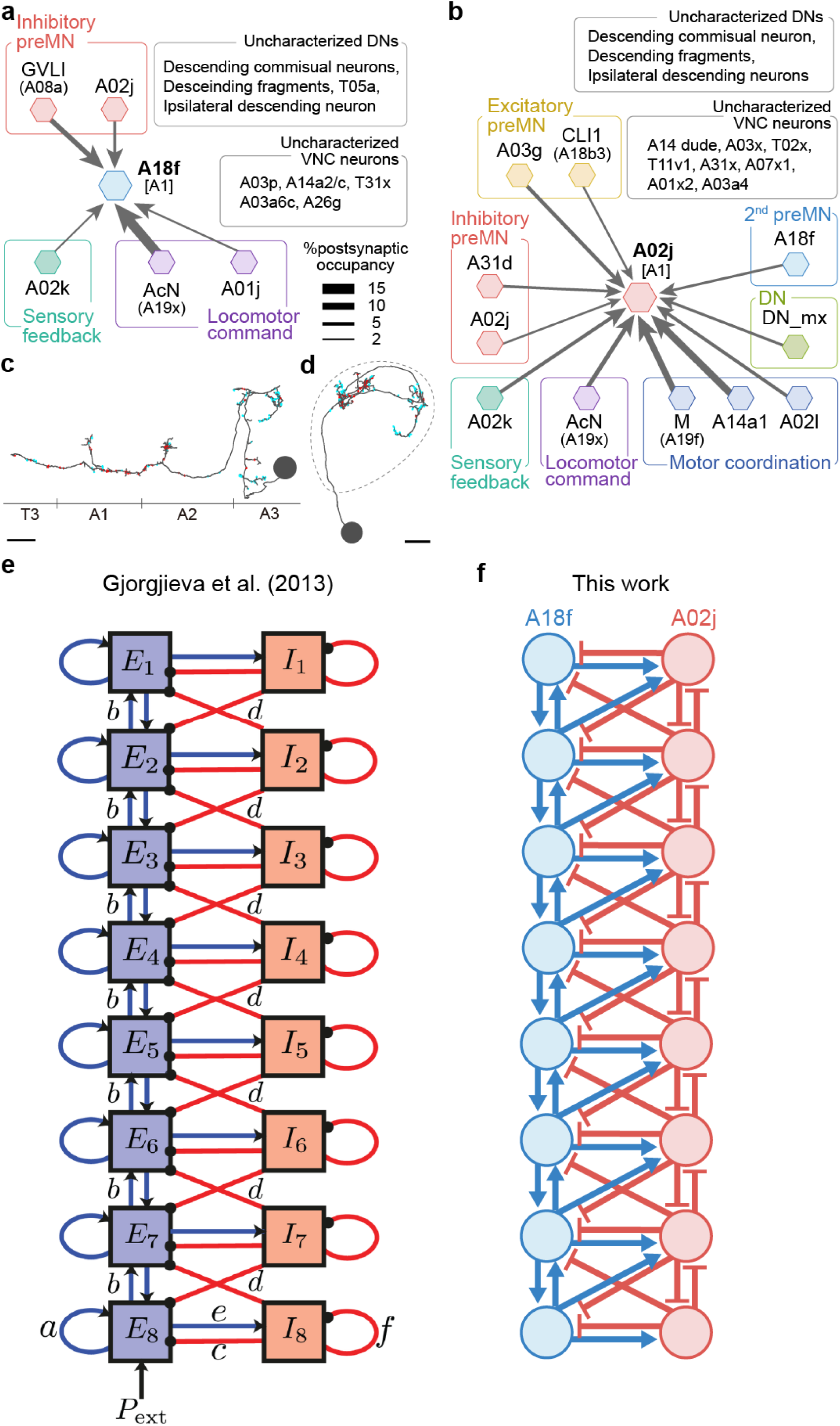
Identification of the A18f-A02j circuit and its architectural similarity to a theoretical model of larval peristalsis. **a**, Identification of A02j neurons as major upstream partners of A18f in the A1 neuromere. **b**, Reciprocal identification of A18f neurons as major upstream partners of A02j in the A1 neuromere. Only neurons with a postsynaptic occupancy of > 2% are shown in **a** and **b**. Arrow thickness is proportional to postsynaptic occupancy. **c, d**, EM reconstruction of A02j neurons. Dorsal view (**c**) and posterior view (**d**) of the A02j neuron in the right A3 hemi-neuromere. Blue and red puncta indicate input and output sites, respectively. The dashed line indicates the boundary of the neuropil. Scale bar, 5 µm. **e**, **f**, The theoretical circuit motif proposed by Gjorgjieva *et al*. (2013) (**e**) and the A18f-A02j circuit motif identified in this study (**f**). The two motifs share a similar segmentally repeated excitatory–inhibitory architecture with local reciprocal interactions and intersegmental coupling along the body axis. Intersegmental connections spanning more than two neuromeres are omitted for clarity in **f**. Reproduced from Gjorgjieva *et al*. (2013), Frontiers in Computational Neuroscience, under the Creative Commons Attribution License (CC BY).

**Extended data Figure 3.**
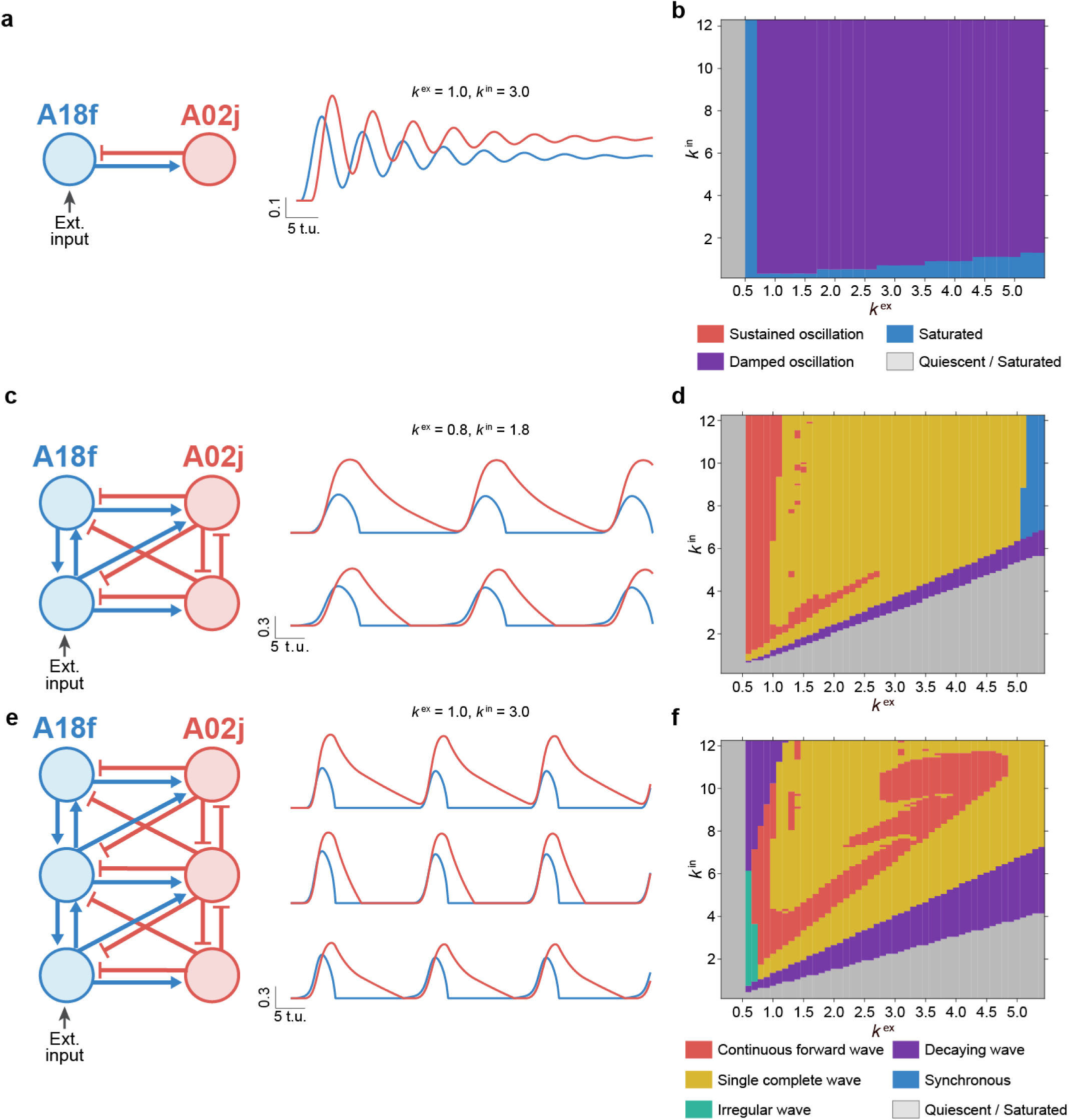
Oscillation and wave generation in truncated A18f-A02j circuit models. **a, b**, Simulation of a single-segment A18f-A02j circuit. **a**, Circuit diagram and representative simulated activity traces showing damped oscillations. *k*^ex^ = 1.0, *k*^in^ = 3.0, external input amplitude = 2.0. **b**, Parameter space analysis. **c, d**, Simulation of a two-segment A18f-A02j circuit. **c**, Circuit diagram and representative simulated activity traces showing propagating activity between the two segments. *k*^ex^ = 0.8, *k*^in^ = 1.8, external input amplitude = 2. **d**, Parameter space analysis. **e, f**, Simulation of a three-segment A18f-A02j circuit. **e**, Circuit diagram and representative simulated activity traces showing forward-propagating waves. *k*^ex^ = 1.0, *k*^in^ = 3.0, external input amplitude = 2. **f**, Parameter space analysis.

**Extended data Figure 4.**
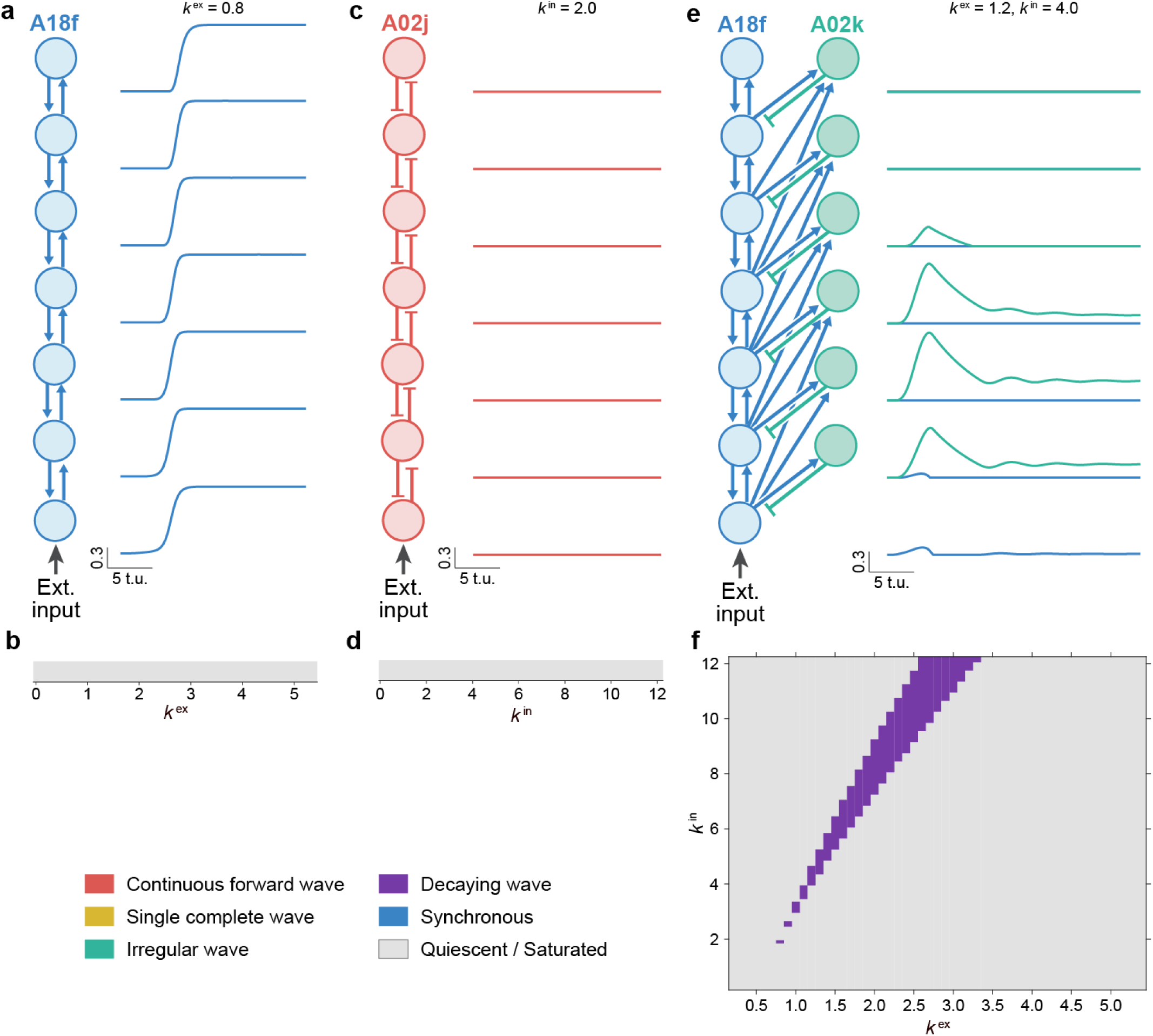
Simulations of control circuits fail to generate forward waves. **a, b,** Simulation of a templated A18f-only circuit. **a**, Circuit diagram and representative simulated activity traces showing failure to generate forward waves. *k*^ex^ = 0.8, external input amplitude = 2.0. **b**, Parameter space analysis. **c, d**, Simulation of a templated A02j-only circuit. **c**, Circuit diagram and representative simulated activity traces showing failure to generate forward waves. *k*^in^ = 2.0, external input amplitude =. **d**, Parameter space analysis. **e, f**, Simulation of a templated A18f-A02k circuit. **e**, Circuit diagram and representative simulated activity traces showing failure to generate forward waves. *k*^ex^ = 1.2, *k*^in^ = 4.0, external input amplitude =. **f**, Parameter space analysis.

**Extended data Figure 5.**
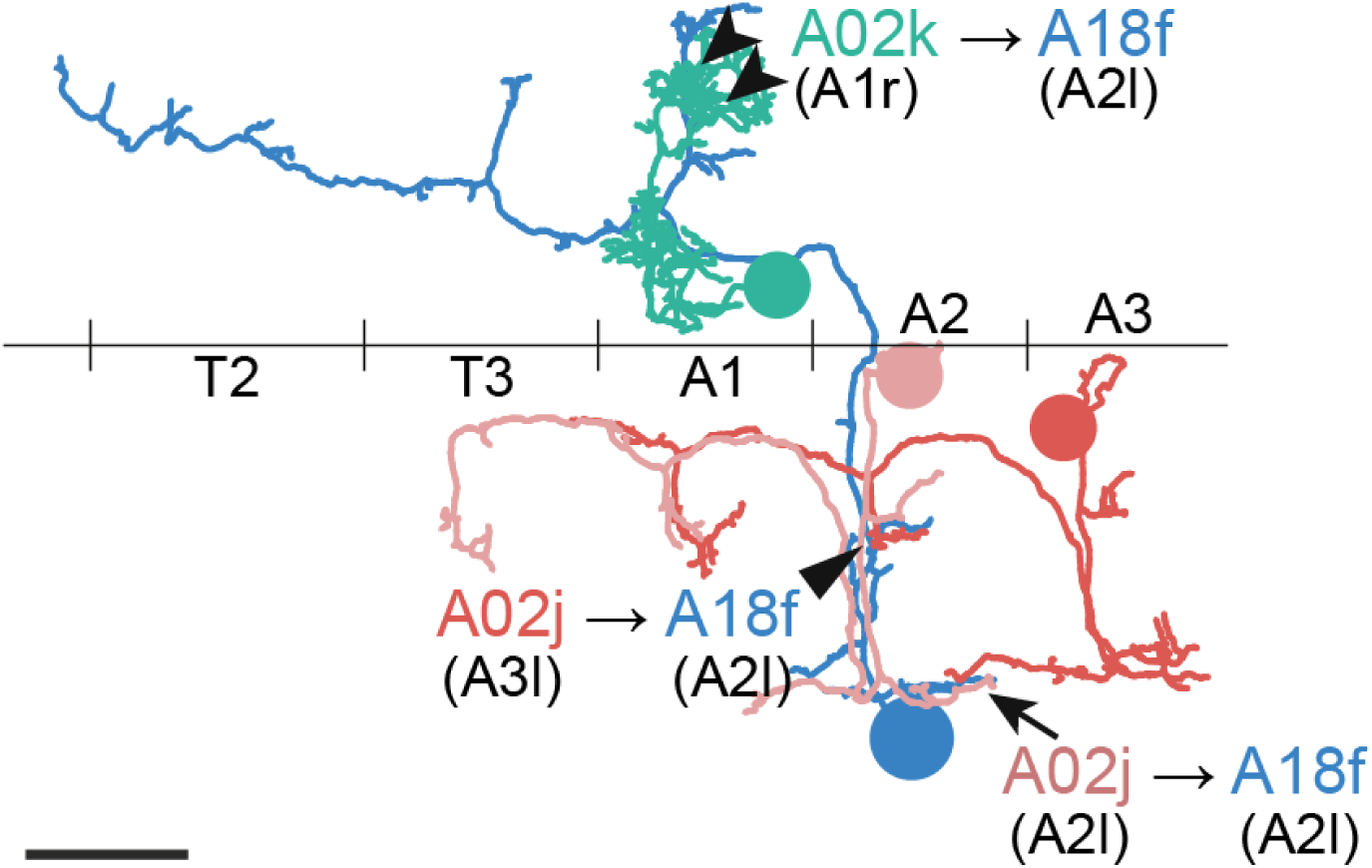
Spatial segregation of A02j and A02k inputs onto A18f neurons. EM reconstruction of input sites onto the A18f neuron in the left A2 hemi-neuromere (blue). Inputs from the same- neuromere A02j neuron (pink) are localized to the proximal dendrite of A18f (arrow), whereas inputs from the posterior A3 A02j neuron (red) are localized to the distal dendrite (triangle). In contrast, inputs from the A02k neuron (green) are localized to collateral branches of the A18f neurite (arrowheads). Scale bar, 10 µm.

**Extended data Figure 6.**
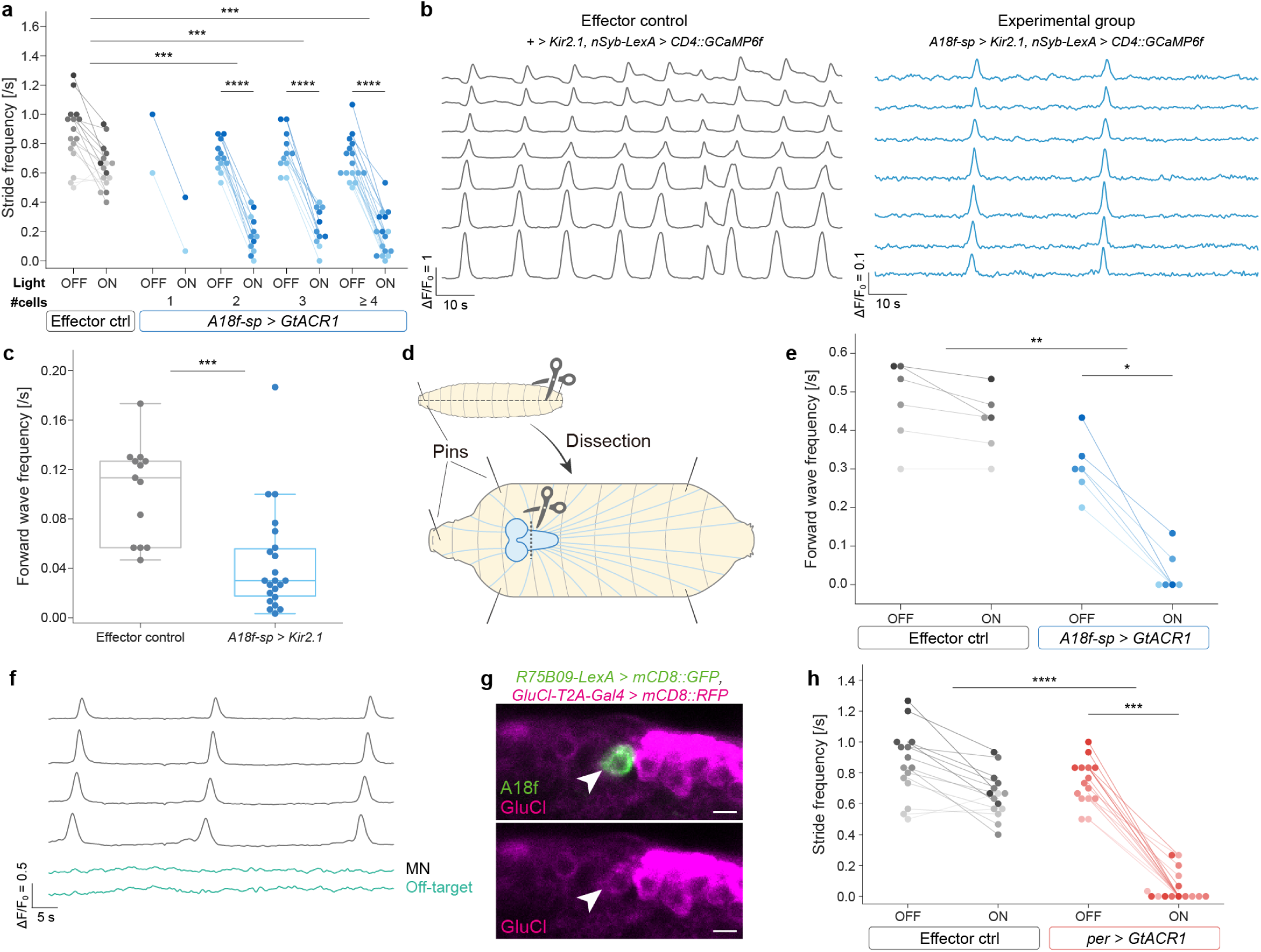
Additional experiments on A18f silencing and glutamatergic inhibition of A18f. **a**, Experimental data from Fig. 5d plotted according to the number of A18f cells expressing GtACR1. Forward locomotion was strongly inhibited even when GtACR1 was expressed in a small number of A18f neurons. *n* = 16, 2, 12, 11 and 16 for effector control, 1, 2, 3, and ≥4 cells, respectively. \*\*\**P* < 0.001, \*\*\*\**P* < 0.0001, paired *t*-test and Welch’s *t*-test with Holm-Bonferroni correction. **b, c,** Chronic silencing of A18f neurons in the isolated CNS reduced the frequency of fictive forward crawling. The CNS was severed posterior to the SEZ, leaving the VNC region containing A18f neurons but excluding the SEZ off- target cells. Pilocarpine, a muscarinic acetylcholine receptor agonist, was applied to increase the frequency of forward crawling. **b**, Representative calcium transients are shown for the effector control (left, *+/UAS-Kir2.1, nSyb-LexA > LexAop-CD4::GCaMP6f* ) and the experimental group (right, *A18f-spGal4 > UAS-Kir2.1, nSyb-LexA > LexAop- CD4::GCaMP6f*). **c**, Quantification of the frequency of fully propagated forward waves. Effector control, *n* = 13; experimental group, *n* = 22. \*\*\**P*<0.001, Welch’s *t*-test. **d, e**, Acute optogenetic silencing of A18f neurons in semi-intact preparations strongly inhibited forward waves. **d**, Schematic of the semi-intact preparation. After dissection, the CNS was severed posterior to the SEZ to exclude off- target cells. **e**, Quantification of forward waves before and during light irradiation in the effector control (grey, *+/UAS- GtACR1*, *n* = 6) and experimental group (blue, *A18f-spGal4 > UAS-GtACR1*, *n* = 6). Optogenetic silencing reduced the frequency of forward waves in the experimental group, indicating that A18f neurons, rather than SEZ off-target cells, are required for forward wave generation. \**P* < 0.05, \*\**P* < 0.01, exact unpaired permutation test and exact paired permutation test with Holm-Bonferroni correction. **f**, Off-target cells labelled by A18f-spGAL4 showed no detectable activity during fictive locomotion. Calcium transients of motor neurons (grey) and an off-target cell (green) are monitored using *A18f-spGal4 > UAS- CD4::GCaMP6f* and *eve-Gal4 > UAS-CD4::GCaMP6f*. **g**, *GluClα* expression in A18f neurons. *GluClα*-positive neurons were labeled by *GluClα-T2A-Gal4 > UAS- mCD8::RFP* (magenta) and A18f neurons are labeled by *R75B09-LexA > LexAop-mCD8::GFP* (green). Arrowhead indicates the soma of an A18f neuron. Scale bar, 5 µm. **h**, Transient silencing using *per-Gal4* driven GtACR1 strongly inhibited larval forward locomotion. Forward locomotion was quantified before and during light irradiation in the effector control (grey, *+/UAS-GtACR1*, *n* = 16) and experimental group (red, *per-Gal4 > UAS-GtACR1*, *n* = 15). \*\*\**P* < 0.001, \*\*\*\**P* < 0.0001, paired permutation Brunner-Munzel test and Welch’s *t*-test with Holm-Bonferroni correction.

**Extended Data Figure 7.**
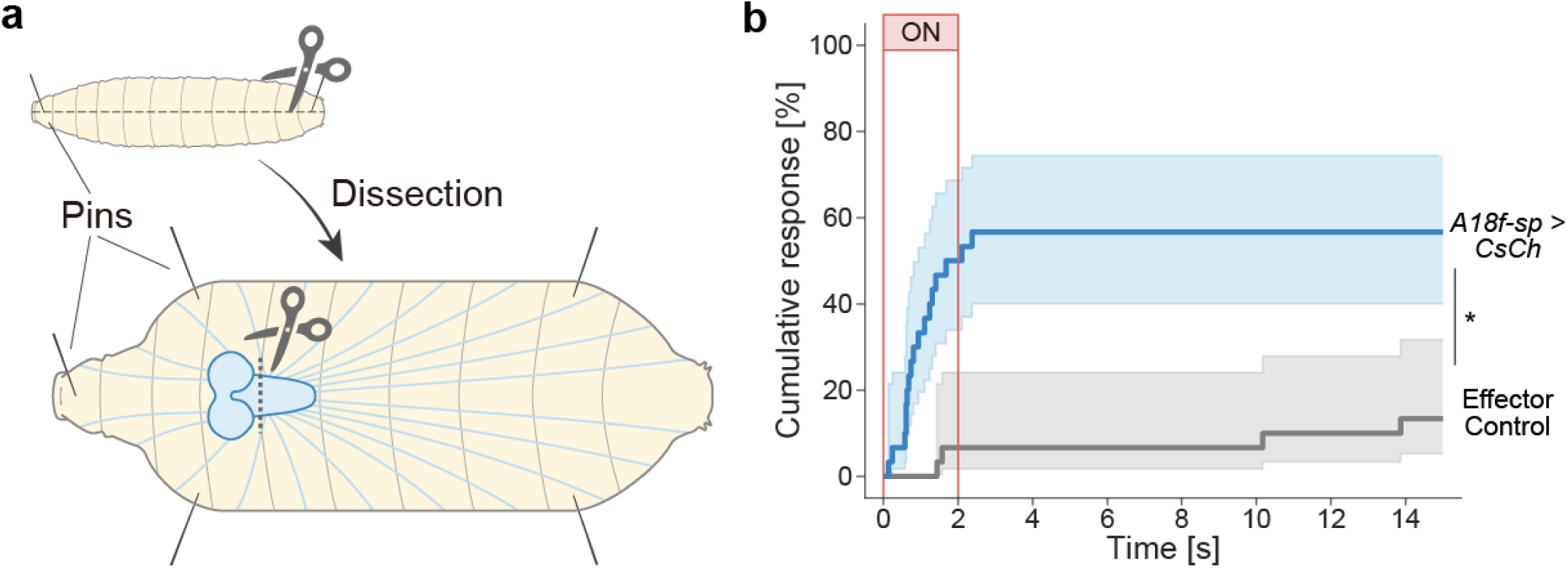
A18f activation induces forward waves in semi-intact preparations lacking the SEZ. **a, b**, Optogenetic activation of A18f-spGal4-labelled neurons induced forward waves in semi-intact preparations after the CNS was severed posterior to the subesophageal zone (SEZ), implicating A18f neurons, rather than SEZ off-target cells, in forward wave induction. **a**, Schematic of the experiment. **b**, Cumulative incidence of forward waves following light onset in semi-intact preparations. Solid lines and shaded regions represent the Kaplan-Meier estimates and 95% confidence intervals, respectively. Effector control, grey, *UAS-CsChrimson/+,* n = 30 trials from 10 animals; experimental group, blue, *A18f-spGal4 > UAS-CsChrimson*, n = 30 trials from 10 animals. \**P* < 0.05, Cox proportional hazards model with cluster-robust standard errors.

