## Supplementary Legends for "A canonical excitatory–inhibitory motif embedded in the connectome underlies rhythmic locomotion in *Drosophila* larvae"

### Supplementary information guide

**Supplementary Note.**

**Mathematical analysis of rhythm generation in the A18f-A02j network.**

This note provides a linear stability analysis of the A18f-A02j network and examine how intersegmental coupling can enable rhythmic activity.

**Supplementary Table 1.**

Downstream targets of A18f neurons identified from connectome data (A2–A4). The downstream neurons are restricted to those located in the A1 neuromere. Only neurons with postsynaptic occupancy greater than 5% are shown.

**Supplementary Table 2.**

Upstream targets of A18f neurons identified from connectome data (A1). Only neurons with postsynaptic occupancy greater than 2% are shown.

**Supplementary Table 3.**

Upstream targets of A02j neurons identified from connectome data (A1). Only neurons with postsynaptic occupancy greater than 2% are shown.

**Supplementary Table 4.**

Connectivity matrix of the A18f/A02j circuit derived from connectome data. Columns represent presynaptic neurons, and rows represent postsynaptic neurons. Left and right homologous neurons within each neuromere are pooled into a single node.

**Supplementary Table 5.**

Connectivity matrix of the A18f/A02k circuit derived from connectome data. Columns represent presynaptic neurons, and rows represent postsynaptic neurons. Left and right homologous neurons within each neuromere are pooled into a single node.

**Supplementary Video 1.**

**Transient silencing of A18f neurons strongly impaired forward locomotion.**

Representative videos of the effector control (*+/UAS-GtACR1*) and the experimental group expressing GtACR1 in A18f neurons (*A18f-spGal4 > UAS-GtACR1*). The video shows a 10-s period of light irradiation, together with the 5-s periods immediately before and after irradiation. The light ON/OFF status is indicated in the top left corner.

The video is shown in real time (1x speed).

**Supplementary Video 2.**
**Chronic silencing of A18f neurons impaired forward locomotion.**
Representative videos of the effector control (*+/UAS-Kir2.1*) and the experimental group expressing Kir2.1 in A18f neurons (*A18f-spGal4 > UAS-Kir2.1*). The video is shown in real time (1x speed).

**Supplementary Video 3.**
**Brief activation of A18f neurons induced forward locomotion in resting larvae.**
Representative videos of the effector control (*+/UAS-CsChrimson*) and the experimental group expressing CsChrimson in A18f neurons (*A18f-spGal4 > UAS-CsChrimson*). The video shows a 1-s period of light irradiation, together with the 10-s periods immediately before and after irradiation. The light ON/OFF status is indicated in the top left corner. The video is shown in real time (1x speed).

**Supplementary Video 4.**
**Brief activation of A18f neurons induced forward waves in semi-intact preparations.**
Representative videos of the effector control (*+/UAS-CsChrimson*) and the experimental group expressing CsChrimson in A18f neurons (*A18f-spGal4 > UAS-CsChrimson*). The larva was pinned flat to a silicone surface as a filleted semi-intact preparation. Anterior is to the left. The video shows a 2-s period of light irradiation, together with the 10-s periods immediately before and after irradiation. The light ON/OFF status is indicated in the top left corner. The video is shown in real time (1x speed).
