## Supplementary Note for "A canonical excitatory–inhibitory motif embedded in the connectome underlies rhythmic locomotion in *Drosophila* larvae"

**Supplementary Note: Linear stability analysis and mode expansion of the A18f-A02j circuit**

In the Wilson–Cowan oscillator, the system exhibits a stable limit cycle under specific parameters. In contrast, a single-segment A18f-A02j loop exhibits decaying oscillations toward a fixed point (Extended data Fig. 3a). This is likely because it lacks recurrent connections from E nodes to E nodes. However, when multiple A18f-A02j loops are connected in a specific wiring pattern, they generate stable oscillations; this phenomenon is also observed in a circuit consisting of two A18f-A02j loops (Extended data Fig. 3c, d). Below, we perform an approximate linear stability analysis to elucidate the origin of these dynamics. To analytically elucidate how network coupling generates oscillatory potential, we consider a simplified, symmetric two-segment approximation. Note that while this linear analysis explains the emergence of the core rhythmic oscillation, the specific phase delays characterizing actual traveling waves arise from structural and boundary asymmetries present in the full simulation.

First, the equation for the firing rate model in this study can be written as

$$\tau \dot{r}_i = f_i(\boldsymbol{r}) \tag{1}$$

where

$$f_i(\boldsymbol{r}) = -V_0 - r_i + I_{\mathrm{ext}} + (r^{\max} - r_i)\sum_j \left(k^{\mathrm{ex}} W^{\mathrm{ex}}_{ij} r_j\right) - \sum_j \left(k^{\mathrm{in}} W^{\mathrm{in}}_{ij} r_j\right) \tag{2}$$

Here, $r_i$ is the firing rate of cell $i$, $V_0$ is the activity threshold, $I_{\mathrm{ext}}$ is the external input, $r^{\max}$ is the maximum firing rate, $W$ is the matrix where the $(i, j)$ component represents the number of synapses connecting cell $j$ to cell $i$, $W^{\mathrm{ex}}$ and $W^{\mathrm{in}}$ denote the excitatory and inhibitory connections extracted from $W$, respectively, and $k^{\mathrm{ex}}$ and $k^{\mathrm{in}}$ are the efficacies of the excitatory and inhibitory synapses, respectively.

Here, we consider excitatory cells $E_1, E_2$ and inhibitory cells $I_1, I_2$. The linearized equation governing the small fluctuation $\Delta r_i(t) = r_i(t) - r_i^*$ around the fixed point $\boldsymbol{r}^*$ can be expressed using the Jacobian matrix $J$ as

$$\tau \Delta \dot{r}_i(t) \simeq \sum_j J_{ij} \Delta r_j \tag{3}$$

Here, $J_{ij} = \partial f_i / \partial r_j |_{\boldsymbol{r}=\boldsymbol{r}^*}$ is calculated as follows.

**(i) For diagonal components ($i = j$)**

Evaluating the derivative at the fixed point $\boldsymbol{r}^*$, and utilizing the fact that there are no direct self-connections in this network architecture ($W_{ii} = 0$), yields:

$$J_{ii} = -1 - \sum_k (k^{\mathrm{ex}} W^{\mathrm{ex}}_{ik} r^*_k) =: -1 - A_i \tag{4}$$

where $A_i \geq 0$ represents an effective self-decay term.

**(ii) For off-diagonal components ($i \neq j$)**

Evaluating the derivative gives:

$$J_{ij} = (r^{\max} - r^*_i)\, k^{\mathrm{ex}} W^{\mathrm{ex}}_{ij} - k^{\mathrm{in}} W^{\mathrm{in}}_{ij} =: B_i k^{\mathrm{ex}} W^{\mathrm{ex}}_{ij} - k^{\mathrm{in}} W^{\mathrm{in}}_{ij} \tag{5}$$

where $B_i \geq 0$ represents the remaining capacity to receive excitation.

Based on these derivatives, we define the effective connection strengths from cell $j$ to cell $i$ as $w_{EE,ij} = B_{E_i} k^{\text{ex}} W^{\text{ex}}_{E_i E_j}$, $w_{EI,ij} = k^{\text{in}} W^{\text{in}}_{E_i I_j}$, $w_{IE,ij} = B_{I_i} k^{\text{ex}} W^{\text{ex}}_{I_i E_j}$, and $w_{II,ij} = k^{\text{in}} W^{\text{in}}_{I_i I_j}$. The small fluctuations of the four cells follow the equations:

$$\begin{aligned}
\tau \Delta \dot{r}_{E_1} &= (-1 - A_{E_1})\Delta r_{E_1} + w_{EE,12}\Delta r_{E_2} - w_{EI,11}\Delta r_{I_1} - w_{EI,12}\Delta r_{I_2} \\
\tau \Delta \dot{r}_{E_2} &= w_{EE,21}\Delta r_{E_1} + (-1 - A_{E_2})\Delta r_{E_2} - w_{EI,21}\Delta r_{I_1} - w_{EI,22}\Delta r_{I_2} \\
\tau \Delta \dot{r}_{I_1} &= w_{IE,11}\Delta r_{E_1} + w_{IE,12}\Delta r_{E_2} + (-1 - A_{I_1})\Delta r_{I_1} - w_{II,12}\Delta r_{I_2} \\
\tau \Delta \dot{r}_{I_2} &= w_{IE,21}\Delta r_{E_1} + w_{IE,22}\Delta r_{E_2} - w_{II,21}\Delta r_{I_1} + (-1 - A_{I_2})\Delta r_{I_2}
\end{aligned} \tag{6}$$

To extract the core mechanism of wave generation, we assume the parameters are symmetric across the two segments ($A_{X_1} = A_{X_2} \equiv A_X$, $w_{XY,11} = w_{XY,22} \equiv w_{XY,\text{intra}}$, and $w_{XY,12} = w_{XY,21} \equiv w_{XY,\text{inter}}$ for $X, Y \in \{E, I\}$). We perform a change of variables into cooperative modes ($E_+ = \Delta r_{E_1} + \Delta r_{E_2}$, $I_+ = \Delta r_{I_1} + \Delta r_{I_2}$) and differential modes ($E_- = \Delta r_{E_1} - \Delta r_{E_2}$, $I_- = \Delta r_{I_1} - \Delta r_{I_2}$).

By adding and subtracting the corresponding equations in (6), the system decouples into the macroscopic dynamics of each mode. Defining the effective macroscopic interactions $w_E = w_{EE,\text{inter}}$, $w_I = w_{II,\text{inter}}$, $w_{EI,+} = w_{EI,\text{intra}} + w_{EI,\text{inter}}$, and $w_{IE,+} = w_{IE,\text{intra}} + w_{IE,\text{inter}}$, the cooperative modes follow:

$$\begin{aligned}
\tau \dot{E}_+ &= (-1 - A_E + w_E)E_+ - w_{EI,+} I_+ \\
\tau \dot{I}_+ &= w_{IE,+} E_+ + (-1 - A_I - w_I) I_+
\end{aligned} \tag{7}$$

Similarly, defining the differential effective interactions $w_{EI,-} = w_{EI,\text{intra}} - w_{EI,\text{inter}}$ and $w_{IE,-} = w_{IE,\text{intra}} - w_{IE,\text{inter}}$, the differential modes follow:

$$\begin{aligned}
\tau \dot{E}_- &= (-1 - A_E - w_E)E_- - w_{EI,-} I_- \\
\tau \dot{I}_- &= w_{IE,-} E_- + (-1 - A_I + w_I) I_-
\end{aligned} \tag{8}$$

Because of the excitatory intersegmental coupling $w_E > 0$, the differential mode $E_-$ experiences a strong decay given by $-1 - A_E - w_E$. Assuming the intrinsic decay $-1 - A_I$ stabilizes the differential mode $I_-$, the fast decay of phase deviations ensures that the long-term dynamics are strictly governed by the cooperative macroscopic modes $(E_+, I_+)$ representing the segmentally coordinated network.

To illustrate the impact of intersegmental coupling on the system's stability, we compare the Jacobian matrix of a single, isolated E-I loop ($J_{\text{single}}$) with that of the dimensionally reduced, coupled macroscopic E-I system ($J_{\text{coupled}}$).

For the single isolated loop, the Jacobian is:

$$J_{\text{single}} = \begin{pmatrix} -1 - A_E & -w_{EI,\text{intra}} \\ w_{IE,\text{intra}} & -1 - A_I \end{pmatrix} \tag{9}$$

Here, the trace is strictly negative ($T_{\text{single}} = -2 - A_E - A_I < 0$) and the determinant is strictly positive ($D_{\text{single}} = (1 + A_E)(1 + A_I) + w_{EI,\text{intra}} w_{IE,\text{intra}} > 0$), resulting in asymptotic stability (decaying oscillations towards a fixed point).

In contrast, the effective Jacobian acting on the macroscopic modes $(E_+, I_+)$ in the coupled system becomes:

$$J_{\text{coupled}} = \begin{pmatrix} -1 - A_E + w_E & -w_{EI,+} \\ w_{IE,+} & -1 - A_I - w_I \end{pmatrix} \tag{10}$$

The reciprocal connections across segments fundamentally alter the diagonal elements of the cooperative mode: the intersegmental E-E connections act as an effective self-excitation $(+w_E)$, while the intersegmental I-I connections act as an effective self-inhibition $(-w_I)$. The trace of this coupled system is:

$$T_{\text{coupled}} = -2 - A_E - A_I + w_E - w_I \tag{11}$$

If the intersegmental excitatory connection strength $w_E$ is sufficiently large to overcome both the intrinsic cellular decay and the intersegmental inhibition $w_I$, the trace turns positive ($T_{\text{coupled}} > 0$). Provided that the determinant of the Jacobian remains positive—a condition generally maintained by the strong reciprocal negative feedback between E and I populations—the real parts of the eigenvalues become positive, destabilizing the fixed point through a Hopf bifurcation.

In conclusion, although a single A18f-A02j loop lacking recurrent connections does not possess a stable limit cycle, the reciprocal excitatory connections across segments act as an effective recurrent excitation-like term. While this linear analysis demonstrates the destabilization of the quiescent state, numerical simulations confirm that the network's nonlinearities restrict this instability, thereby generating the stable, sustained oscillations required for locomotor rhythms.
